# A simulation-based method for genotype-environment association analysis

**DOI:** 10.64898/2026.08.23.746561

**Authors:** Takahiro Sakamoto, Sam Yeaman

**Affiliations:** Department of Biological Sciences, University of Calgary, Calgary, AB T2N 1N4, Canada; National Institute of Genetics, Mishima, Shizuoka 411-8540, Japan; Department of Biology, Faculty of Science, Kyushu University, Fukuoka 819-0395, Japan

**Keywords:** local adaptation, genotype-environment association, statistical method, coalescent theory

## Abstract

Genotype-environment association (GEA) analyses are widely used to identify loci underlying local adaptation by examining correlations between allele frequencies and environmental variables across a species’ range. A major challenge for this approach is distinguishing true adaptive signals from spurious associations arising from population structure. Several methods have been developed to account for population structure, but these methods can suffer from reduced statistical power or increased false positives under some conditions. To address this, we introduce a new GEA method, termed *SimGEA*. In essence, SimGEA infers a neutral evolutionary model that reproduces the population structure observed in empirical data and uses this model to simulate neutral alleles. By applying the same GEA statistic to both the empirical and simulated data, SimGEA evaluates the significance of observed associations against neutral expectations that account for population structure. We compared the performance of SimGEA with that of existing GEA methods, including LFMM2 and BayPass, using simulations of local adaptation in two-dimensional space. We found that SimGEA consistently controlled the false discovery rate without substantially sacrificing statistical power across the scenarios examined. These results suggest that calibrating statistics using neutral simulations provides a robust and flexible approach for accounting for population structure in GEA analyses.

---

Local adaptation is widely observed among species inhabiting heterogeneous environments (Felsenstein 1976; Hereford 2009; Wadgymar *et al*. 2022). Elucidating its genetic basis is fundamental to understanding how populations respond to spatial environmental variation through evolution (Savolainen *et al*. 2013; Booker *et al*. 2026). However, identifying genes underlying local adaptation is often challenging because signals of adaptation are confounded with population structure shaped by random genetic drift.

A widely used signal for detecting genomic regions involved in local adaptation is the correlation between allele frequencies and environmental variables within a species (Lasky *et al*. 2023). When different alleles are favored and reach high frequency in different environments, these correlations can become pronounced (Figure 1a). Accordingly, many studies have used correlation or regression tests to identify outlier variants showing genotype-environment associations (Joost *et al*. 2007; Poncet *et al*. 2010; Jones *et al*. 2013). Of note, redundancy analysis (RDA) is a powerful approach when multiple environmental variables are to be tested (Forester *et al*. 2016, 2018; Capblancq *et al*. 2018). However, a major drawback for these approaches is their failure to account for population structure (Meirmans 2012; De Mita *et al*. 2013). In spatially structured populations, isolation by distance can generate allele-frequency clines across habitats. When environmental variables also vary gradually in space, such as north-south gradient of temperature, significant correlations can arise even at neutral loci (Figure 1b), complicating their interpretation. Therefore, although these simple association tests may still be useful for identifying candidate outlier loci (Yeaman *et al*. 2016; Forester *et al*. 2018; Capblancq *et al*. 2023), explicitly accounting for population structure is desirable (Hoban *et al*. 2016).

**Figure 1.**
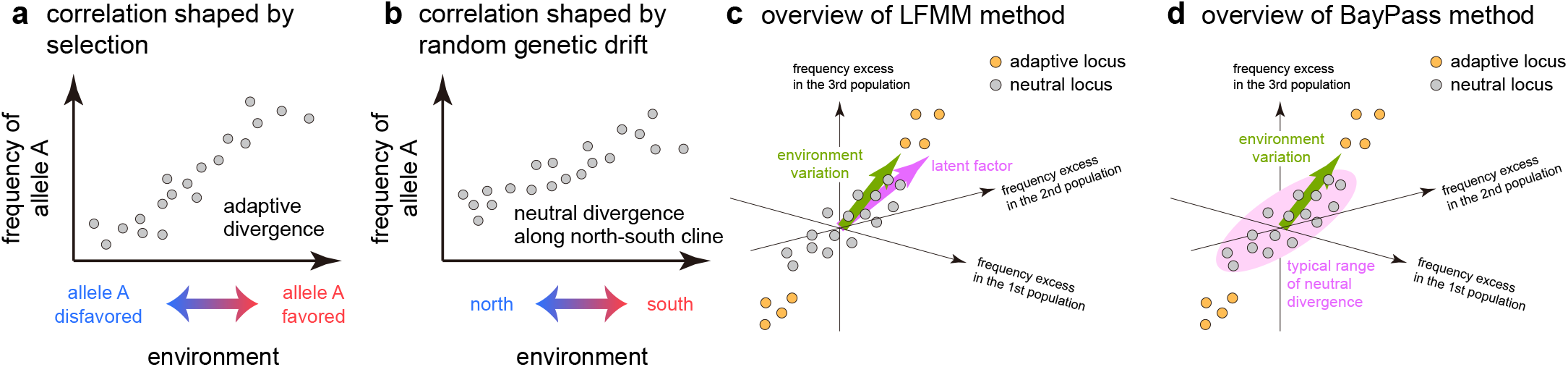
Conceptual overview of genotype-environment association (GEA) analysis. (a) A strong correlation between genotype and an environmental variable can arise when a focal locus is subject to environment-dependent selection (i.e., divergent selection). (b) Similar correlations can also be generated by neutral processes such as isolation by distance, particularly when environmental variables exhibit gradient spatial patterns. (c) Schematic illustration of how LFMM accounts for population structure in GEA analyses. Latent factors (pink arrow) are estimated to capture major axes of neutral allele-frequency variation. Genotype-environment associations are then tested by regression on the environmental variable (green arrow) while including these latent factors as covariates. Therefore, it is expected that the regression does not work well when the focal environment well aligns with one of the latent factors. (d) Schematic illustration of how BayPass (and BayEnv) accounts for population structure. The typical range of neutral allele-frequency divergence (i.e., the covariance structure among populations) is modeled using a multivariate normal distribution. This neutral model is compared with the model including environmental variables to assess whether environmental variation helps explain the observed allele-frequency divergence at each locus.

To date, two major approaches have been widely used to explicitly account for population structure in genotype-environment association (GEA) analyses. The first is LFMM (latent factor mixed models), a regression-based framework that incorporates latent factors capturing neutral patterns of allele-frequency variation across space (Frichot *et al*. 2013; Caye *et al*. 2019) (Figure 1c). By including the latent factors as covariates, LFMM detects statistically significant genotype-environment associations while controlling for population structure. The second is BayPass (Gautier 2015), an MCMC-based Bayesian framework that extends BayEnv (Coop *et al*. 2010; Günther and Coop 2013) (Figure 1d). In this framework, the covariance of allele frequencies among subpopulations, which summarizes the effect of population structure, is modeled by a multivariate normal distribution. This neutral covariance model is then compared with an adaptive model including environmental effects to identify loci that deviate significantly from neutral expectations. In addition to these methods, RDA also allows for explicit population structure correction in a similar manner as a partial regression; however, Forester *et al*. (2018) reported that it leads to substantial loss of power.

Although LFMM and BayPass often outperform simple correlation/regression analyses without the population structure correction (de Villemereuil *et al*. 2014), several studies have noted potential limitations of these methods. For LFMM, statistical power may be decreased when the focal environmental variable exhibits a gradual spatial pattern (Lotterhos and Whitlock 2015). In such cases, latent factors may align with the environmental gradient, reducing the power of the partial regression and potentially causing truly adaptive signals to remain undetected (Figure 1c). BayPass may be more robust to this issue because it can evaluate excess divergence even when the directions of adaptive and neutral divergence are highly correlated (Figure 1d). However, because BayPass relies on MCMC-based inference, its analytical outcomes may be sensitive to MCMC convergence properties (Blair *et al*. 2014). Furthermore, although both methods are expected to perform well when allele-frequency divergence approximately follows a normal distribution, this assumption may break down when one allele is rare. Under strong spatial structure, local allele frequencies can take extreme values even when their global mean is intermediate, undermining the validity of normality assumptions. Concordantly, some studies used simulated data to evaluate their performance and reported that these methods can show the elevated false-discovery rates under some scenarios (de Villemereuil *et al*. 2014; Rellstab *et al*. 2015; Forester *et al*. 2016; Booker *et al*. 2024).

Recently, Goel *et al*. (2025, 2026) developed a Bayesian framework that aims to improve these methods by partially incorporating an evolutionary mechanistic model. However, its performance has not yet been comprehensively compared against existing GEA methods under diverse demographic scenarios, possibly owing to its high computational cost.

In this study, we propose a new simulation-based method for GEA analysis, *SimGEA* (Figure 2), to address these limitations. The basic philosophy of our approach is similar to that of BayPass, but it does not rely on MCMC algorithms or normal-distribution approximations. First, we estimate the covariance structure of allele frequencies across subpopulations using an analytical approximation rather than MCMC. We then infer a migration matrix that reproduces this covariance structure, using coalescent theory combined with an optimization procedure. The goal of this step is not to estimate the true demographic history, but to construct a model that captures the covariance structure observed in the data (see Methods). This model is subsequently used to perform neutral simulations, generating hypothetical datasets of neutral alleles. These simulated data are used to calculate null distributions for the chosen test statistic, allowing us to evaluate the significance of observed genotype-environment associations while controlling for population structure. Although SimGEA is slower than LFMM2, it can handle *∼* 100 populations within feasible computational time.

**Figure 2.**
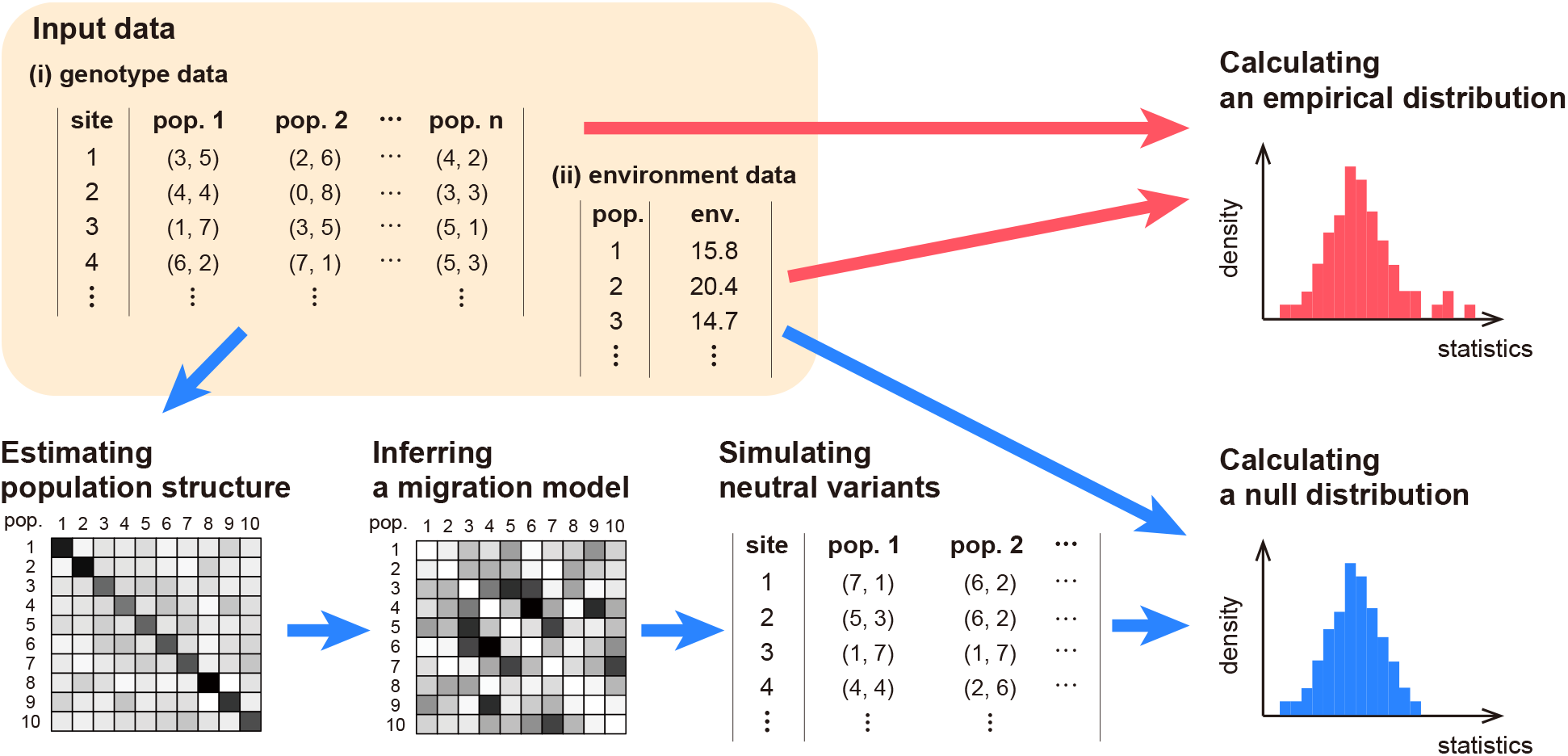
Overview of the SimGEA method. SimGEA requires two input data: (i) allele counts for each subpopulation (i.e., sampling location) at each SNP site, and (ii) environmental data for each subpopulation. Our method first uses the genotype data and estimates a normalized covariance matrix of allele frequency, **Λ** (Step 1). Next, a migration parameter that reproduce the estimated covariance matrix is inferred (Step 2). Using this migration rate, neutral alleles are simulated (Step 3). Ideally, these neutral alleles have the same population structure as the input data while free from the effect of selection. Users then apply the same GEA analysis to the input genotype data and the simulated data using the input environmental data to evaluate the significance of association in the input data.

In the following sections, we first present the proposed method. Next, to evaluate its utility, we compare the performance of previous methods and SimGEA (our method), using simulation data generated by two-dimensional spatial models. Our results show that SimGEA exhibits comparable or superior performance across a wide range of parameter spaces, especially in reducing false-positive cases. These findings suggest that SimGEA provides a robust alternative for detecting genotype-environment associations under wide ranges of demographic and environmental conditions.

## Methods

In this section, we introduce our new method for GEA analysis. We assume that samples are collected from *n* subpopulations, and denote by *a*_*i*_ the number of sampled haplotypes in the *i* th subpopulation. The input genotype data consist of allele counts for *L* biallelic loci across these subpopulations. To minimize the effect of genotyping errors, users may optionally apply a minor allele frequency (MAF) filter before running the analysis.

For each locus, we denote by *b*_*ij*_ the count of a focal allele at locus *i* in subpopulation *j*. The choice of which allele is treated as the focal allele is arbitrary and does not affect the results. Our current implementation assumes no missing values in the genetic data. Practically, missing data may be imputed using imputation tools, for example by using *LEA* package in R (Frichot and François 2015). The environmental value for the *i*th environmental variable in the *j*th subpopulation is denoted by *e*_*ij*_.

Our approach consists of following three steps (Figure 2):

- Estimating the covariance matrix of allele frequencies as a summary of population structure in the dataset
- Inferring a migration matrix that reproduces the estimated covariance structure
- Generating a null distribution of the test statistic using neutral simulations

In the following, we provide an overview of each step.

### (Step 1) Estimating the covariance matrix of allele frequencies

Our goal in this step is to obtain a consistent estimator of the population covariance structure, while accounting for noise arising from finite sample sizes. To this end, we first consider a model underlying the allele frequency divergence among sub-populations. Let ***p***_*i*_ = (*p*_*i*1_, *p*_*i*2_, *· · ·*, *p*_*in*_)^*T*^ be the vector of allele frequencies across subpopulations at locus *i*. Motivated by previous studies (Pickrell and Pritchard 2012; Bradburd *et al*. 2016), we assume that ***p***_*i*_ can be regarded as random variables that diverge around a global mean frequency:

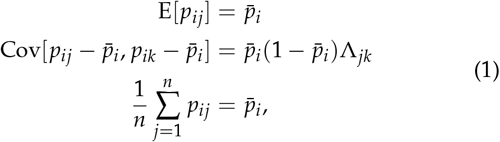

where **Λ** is a *n × n* matrix shared by all neutral loci, and 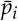 is a locus-specific parameter representing a global mean allele frequency. Here, **Λ** represents the correlated allele-frequency deviations from the global mean shaped by population structure and is assumed not to depend on 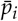. By construction, each row and column of **Λ** sums to zero. Our specific aim of this step is to estimate **Λ** from a sampled genotypic data.

Based on this formulation, the effect of finite sample sizes is considered. We assume that the observed allele count follows a binomial sampling, *b*_*ij*_ *∼* Binom(*a*_*j*_, *p*_*ij*_). Let *z*_*ij*_ = *b*_*ij*_ /*a*_*j*_ be the observed allele frequency in the sample, and denote by 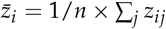 the mean allele frequency calculated from the sample. In Supplementary Text A, we show that *z*_*ij*_ satisfies:

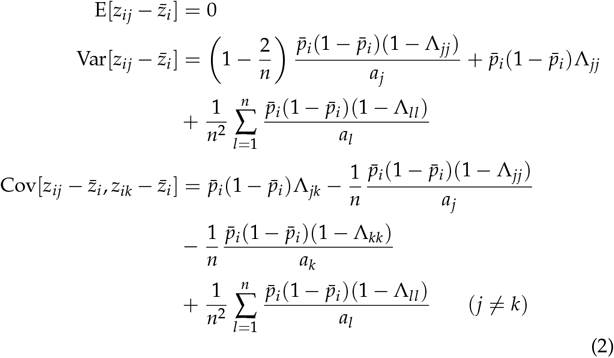

Equation 2 describes the relationship between the sampled allele frequency and the covariance structure. To estimate **Λ** from *z*_*ij*_s, we further assume that the global mean frequency is approximated by the mean value calculated from the sample 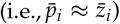. By letting 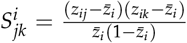, Equation 2 is arranged into

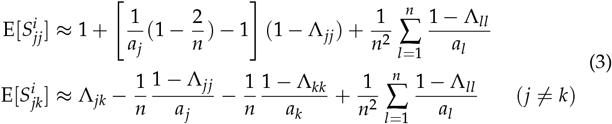

Now, the right hands of Equation 3 do not depend on locus-specific terms. This allows us to estimate the left hand of Equation 3 by the sample average over all loci:

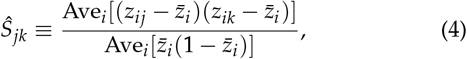

which is the average of observed 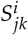 over *L* loci weighted by 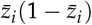. We used the weighted average because rare alleles may be strongly affected by stochasticity and have less reliable information on population structure. These *Ŝ*_*jk*_ s are substituted into the left-side of Equation 3, then this linear equation is numerically solved to calculate **Λ** (Supplementary Text A). We below denote this estimated matrix by **Λ**_obs_.

Although **Λ** is theoretically a positive semi-definite matrix, the estimated matrix **Λ**_obs_ may not satisfy this condition due to the sampling noise. To correct for this, we modify the estimated **Λ**_obs_ by replacing negative eigenvalues with zero if it has negative eigenvalues.

### (Step 2) Inferring a migration matrix

Next, we infer a spatial population model that can reproduce the estimated covariance matrix **Λ** under migration-drift equilibrium. Because **Λ** contains 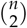 free parameters, a demographic model with a larger number of parameters would result in an identifiability problem. To avoid this, we restrict the search to a model with exactly the same degrees of freedom: *n* subpopulations with homogeneous haploid subpopulation sizes, *N*, and symmetric pairwise migration rates, *m*_*ij*_ = *m*_*ji*_ . Under this parameterization, the equilibrium state is fully characterized by 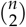 *N*-scaled migration rates, *M*_*ij*_ = 2*Nm*_*ij*_. Assuming a structured coalescent model, the expected covariance can be numerically calculated as a function of ***M*** = (*M*_*ij*_) as **Λ**_theor_(***M***) (see Supplementary Text B for details). Next, we perform an optimization search to find the migration matrix ***M*** that minimizes the discrepancy between the observed covariance matrix **Λ**_obs_ and the theoretical covariance **Λ**_theor_(***M***). For this purpose, we adopt the AdamW algorithm (Loshchilov and Hutter 2017) and define the loss function as *D*(**Λ**_obs_, **Λ**_theor_(***M***)), where *D*(***X, Y***) = ∑_*i<j*_ (*x*_*ij*_ *− y*_*ij*_)^2^. In each optimization step, we need the gradient of this loss function with respect to each migration rate *M*_*ij*_. A straightforward approach would be to numerically compute **Λ**_theor_(***M*** + Δ_*ij*_) where Δ_*ij*_ is a small perturbation to *M*_*ij*_. However, this approach is computationally infeasible when *n* is large, because evaluating **Λ**_theor_(***M***) is computationally demanding, and repeating this computation for all *n*(*n −* 1)/2 entries of ***M*** in every step would incur a prohibitive cost.

To make the optimization scalable, we instead develop a pseudo-gradient that requires computation of **Λ**_theor_(***M***) only once per step. This idea is based on a simple biological intuition: as *M*_*ij*_ increases, allele frequencies in subpopulations *i* and *j* becomes more similar. We assume the effect of a small increase in *M*_*ij*_ as approximately replacing *p*_*i*_ with (1 *− ε*_*ij*_)*p*_*i*_ + *ε*_*ij*_ *p*_*j*_ and *p*_*j*_ with (1 *− ε*_*ij*_)*p*_*j*_ + *ε*_*ij*_ *p*_*i*_ . Using this, change in each element of **Λ**_theor_ may be approximated by

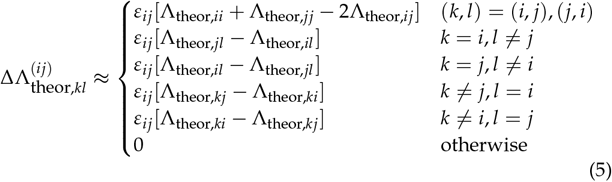

Summing over these terms, the effect of *ε*_*ij*_ on the change in distance is approximated by

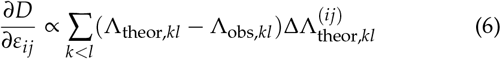

This expression is used as a gradient in our AdamW framework instead of the exact gradient. Please refer to Supplementary Text B for details.

This pseudo-gradient is obviously not an exact derivative, because it does not explicitly model how *ε*_*ij*_ corresponds to a change in *M*_*ij*_ at the migration-drift equilibrium, nor how *M*_*ij*_ affects *p*_*k*_ for *k* ∉ {*i, j*} . However, it provides a rough descent direction while dramatically reducing computational cost. As shown in the Results section, this pseudo-gradient works well in practice.

### (Step 3) Generating a null distribution of the test statistic

Using the migration matrix estimated in Step 2, we generate allele count data of neutral alleles under a structured coalescent model. These simulated datasets are used to construct null distributions of the chosen test statistic, reflecting population structure.

A key note here is that our migration matrix is inferred solely from the covariance structure of the data. As a consequence, the inferred model does not reproduce the allele frequency spectrum in the empirical data. Because many GEA statistics are sensitive to minor allele frequency, the lack of control for it would distort the null distribution. To address this issue, we compared each empirical SNP with simulated alleles with a similar minor allele frequency (Supplementary Text D for details). The choice of test statistic is arbitrary in our framework. The same statistical test is applied to empirical and simulated data, and each SNP is compared against the null distribution corresponding to its frequency.

## Results

In this section, we evaluated the performance of SimGEA in comparison with previous methods using four simulated datasets (Supplementary Text C).

### Accuracy of matrix estimation

As the validity of our method depends on the accuracy of population structure estimation, we first assessed SimGEA’s ability to estimate a migration matrix from genome data. For this purpose, we focused on an island model which falls within the exploration target of inference Step 2 (i.e., uniform *N* and symmetric migration) and examined whether our method can recover the true migration model from the allele count data under this ideal situation. We considered three number of subpopulations (*n* = 10, 30, 100) and three migration strengths (*M*_tot_ = 1 (weak), 10 (intermediate), and 100 (strong)). For each parameter combination, we ran 20 independent simulations to obtain allele counts of 100,000 neutral loci with 8 haploids from each subpopulation, where migration matrices were randomly generated assuming each *M*_*ij*_ *∼* Exponential(*M*_tot_/*n*). Weak, intermediate, strong migration roughly corresponding to pairwise *F*_*ST*_ *≈* 0.35, 0.05, and 0.005, respectively (Figure S1).

To quantify performance, we first examined the correlation between the true and estimated pairwise migration rates (i.e., *M*_*ij*_s) for each replicate (Figure 3a, see Figure S2a for a direct comparison of *M*_*ij*_s). We observed almost perfect correlations for up to *n* = 30 under low to intermediate migration rates, validating our method. The correlation decreased when the number of subpopulations is large (i.e., *n* = 100), or migration is strong such that population structure becomes subtle. This result highlights the difficulty of accurately inferring a migration matrix in a large metapopulation with weak population structure.

**Figure 3.**
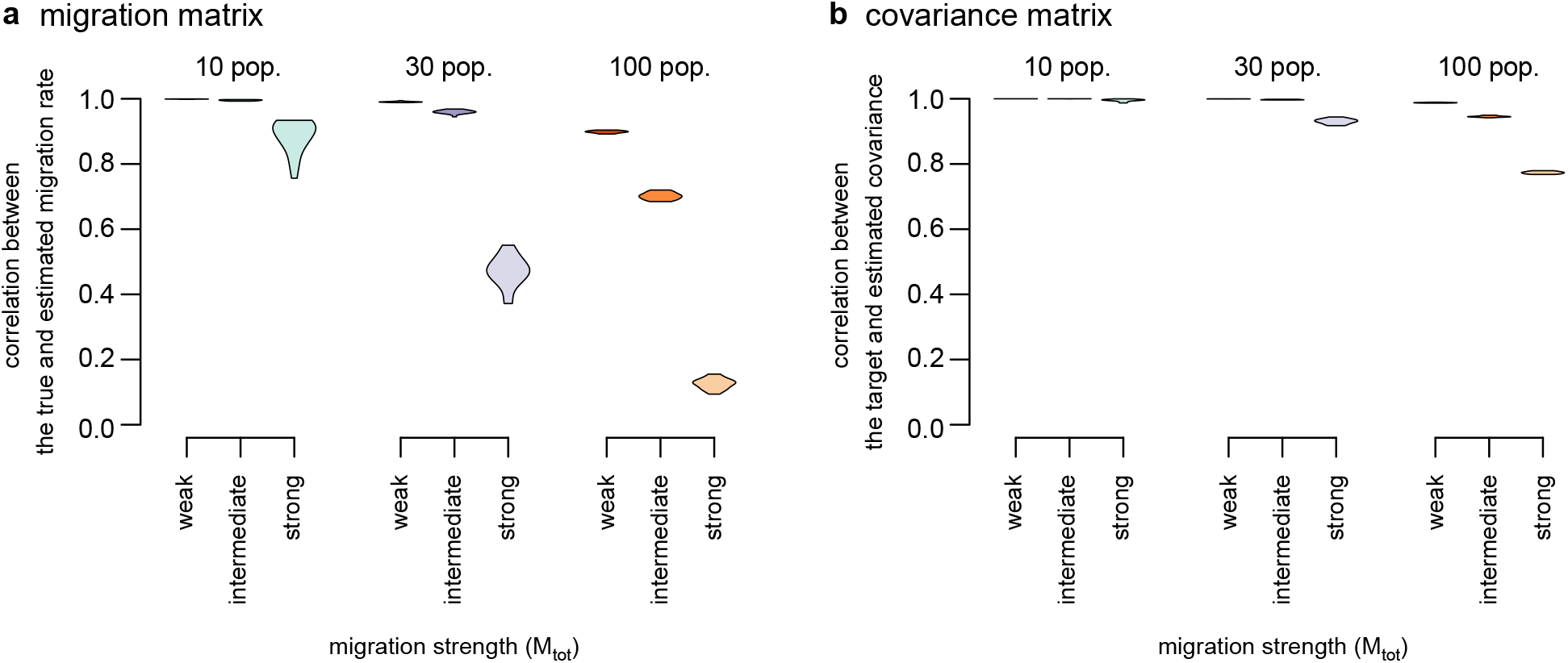
Performance of the estimation of population structure in island models. (a) The distribution of correlation coefficient between the true and estimated migration matrices (*m*_*ij*_ s) in 20 replicates. (b) The distribution of correlation coefficient for covariance matrices (Λ_*ij*_*s*) in 20 replicates.

Despite relatively poor performance for large *n* and large *M*_*ij*_, because our primary focus is on a covariance structure, we also evaluated the performance at the level of covariance structure by comparing **Λ**_theor_(***M***) with **Λ**_obs_ (Figure 3b, see also Figure S2b). Strikingly, our method showed good performance at the covariance level across parameter settings. Notably, the estimates are accurate even in cases with a large metapopulation and/or strong migration, where the estimated migration rate deviates from the true values. This discrepancy may partly arise because the multiple migration models result in similar covariance structures, which may be more likely under a large metapopulation and/or subtle population structure. This pattern suggests that our estimated migration model is useful as a control of population structure although the estimated model does not necessarily reflect the real demography.

Because the framework proposed by Goel *et al*. (2026) also includes an explicit migration-estimation component, we implemented a simplified version of this component for comparison with SimGEA. The original framework can accommodate genotype uncertainty in low-coverage sequencing data, whereas our simulation data consist of error-free allele counts. We therefore omitted the genotype-likelihood data model and focused on the component that estimates a migration matrix from genetic dissimilarities using coalescent theory (see Supplementary Text D). Due to the high computational cost of this Bayesian approach, we analyzed only cases with *n* = 10 and 30.

We found that SimGEA provided more accurate migrationrate estimates for *n* = 10, whereas performance was comparable for *n* = 30 (Figure S3a). Because SimGEA was faster by orders of magnitude (Figure S3b), these results suggest that SimGEA provides a practical and effective option for estimating population structure in this setting. It should be noted, however, that the full framework of Goel *et al*. (2026) explicitly accounts for genotype uncertainty, which is not considered in SimGEA. Therefore, their method may be more appropriate for low-coverage or otherwise error-prone sequencing datasets.

### Performance in two-dimensional space with linkage equilibrium

Next, we evaluated the power to detect loci involving local adaptation under realistic spatial structure. To this end, we considered a two-dimensional 14*×* 14 stepping-stone model (see Supplementary Text C for details). Local adaptation was simulated under a Wright–Fisher model using a single environmental variable with one of the three spatial maps showing different degrees of spatial autocorrelation (Figure 4a). Each map is normalized such that mean and variance of the environmental value across space is 0 and 1, respectively. We assumed two migration rates between adjacent subpopulations (*m* = 0.04 and *m* = 0.01). For *m* = 0.04, average pairwise *F*_*ST*_ between different subpopulations is *≈* 0.024 and the maximum *F*_*ST*_ between the most distant subpopulations is *≈* 0.06, whereas these values are 0.086 and 0.2 for *m* = 0.01, respectively (Figure S4a).

**Figure 4.**
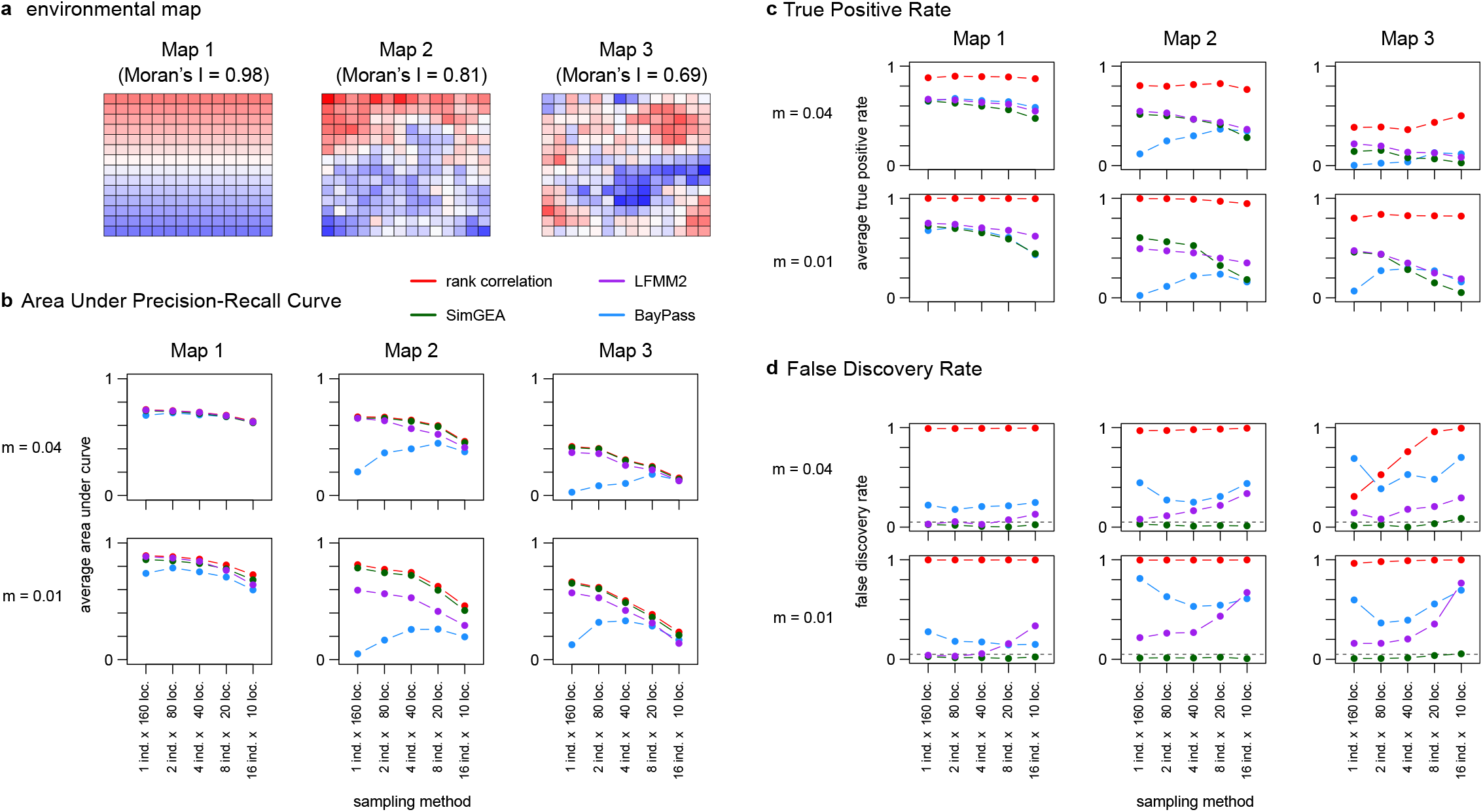
Performance in the two-dimensional stepping stone model with linkage equilibrium. (a) The environmental map assumed in this setting. Each map has different degrees of spatial autocorrelation, represented by Moran’s I of 0.98, 0.81, and 0.69, respectively. (b) The area under the precision-recall curve averaged over 50 replicates. (c) The true positive rate averaged over 50 replicates. (d) The false discovery rate averaged over 50 replicates. The horizontal dashed line represents 0.05, which is the target in the Benjamini–Hochberg procedure.

In addition to neutral loci, we included 15 loci under divergent selection, consisting of five loci each with strong (*s*_*E*_ = 0.01), intermediate (*s*_*E*_ = 0.003), and weak selection (*s*_*E*_ = 0.001). The selection strength at each subpopulation is determined by the product of *s*_*E*_ and the environmental value. During the simulations, linkage equilibrium was assumed among loci, and 160 individuals were sampled at equilibrium. To assess the effect of sampling design, we considered five sampling schemes: sampling *a* = 1, 2, 4, 8, or 16 individuals from *n* = 160, 80, 40, 20, or 10 randomly chosen subpopulations, respectively. After applying a minor allele frequency filter (MAF *≥* 0.05), each dataset typically has 12,000-14,000 neutral polymorphic sites and 10-15 selected polymorphic sites (Figure S4b, c).

Adaptive loci were then inferred using four methods: (i) rank correlation tests without structure correction (Wendt 1972), (ii) LFMM2 (Caye *et al*. 2019), (iii) BayPass (Gautier 2015), and (iv) rank correlation tests calibrated using SimGEA. In SimGEA, a tail approximation based on the generalized Pareto distribution was employed to evaluate significance of outliers accurately from a limited number of simulated variants (Knijnenburg *et al*. 2009, see Supplementary Text D). Overall, this approach produced reasonably well-calibrated p-value estimates across the conditions examined (Supplementary Text E). We did not run the method of Goel *et al*. (2026) due to the limit of computational resources. We ran 50 simulation replicates for each of six parameter combinations (three environmental maps and two migration rates) and evaluated their overall performance.

We first examined whether truly adaptive loci tend to be assigned near the top of the significance rank produced by each method. To quantify this, we computed the area under the precision–recall curve (AUPRC), which is high when adaptive loci are consistently assigned higher ranks than neutral loci.

Across parameters, the simple correlation approach and SimGEA consistently show good performance (Figure 4b). When the environmental variable follows a smooth spatial gradient (Map 1), AUPRC values were similar across methods, although BayPass showed slightly lower performance. In contrast, under partially gradual environments (Maps 2 and 3), BayPass exhibited substantially lower AUPRC when the samples are taken from many locations (*n* ≥ 40), and LFMM2 performed slightly worse than both the simple correlation approach and SimGEA. The performance of the simple correlation and SimGEA was highly similar, reflecting that SimGEA calibrates *p*-values with largely conserving the rank order of loci by design. In general, sampling a larger number of locations with fewer individuals per location resulted in better performance across parameter settings, consistent with Lotterhos and Whitlock (2015).

We also examined the reliability of detected significant associations under a stringent criterion. For this purpose, except for BayPass, loci were defined as significant if their *p*-values were less than 0.05 after correction for multiple testing using the Benjamini–Hochberg procedure. For BayPass, loci were defined as significant if their Bayes factors exceeded 100, following Jeffrey’s rule, without correction for multiple tests. We then computed a true positive rate (TPR) and a false discovery rate (FDR) for each method.

SimGEA achieves effective control of the FDR without substantially sacrificing TPR. Across parameter settings, SimGEA was the only method for which FDRs were consistently around or below the assumed target of 0.05. The simple correlation approach exhibited very high TPR, indicating that a large fraction of truly adaptive loci were identified as significant; however, it also showed elevated FDR due to the lack of correction for population structure. LFMM2 tended to achieve slightly higher power than SimGEA, but this gain was accompanied by increased FDR. LFMM2 shows high FDR when the number of sampling locations are small and/or the environment is not completely gradient (i.e., Maps 2 and 3). BayPass showed limited performance in terms of both TPR and FDR, consistent with the pattern in AUPRC.

In addition to the statistical performance, we also measured the computational time required for each method (Figure S5). LFMM2 was the fastest method, with the regression step taking less than one minute once the number of latent factors was specified. The computational time for SimGEA and BayPass increased with the number of subpopulations (i.e., the number of sampling locations). Within the parameter range examined here, SimGEA tended to be faster than BayPass, although computation times were comparable for some parameter settings. Both methods completed within a half day for all of our datasets.

### Performance under linkage disequilibrium

We next relaxed the assumption of linkage equilibrium, as linkage can confound many genome scan methods (Lotterhos 2019). While the overall simulation framework remained the same as in the linkage equilibrium scenarios, we explicitly modeled genome structure (see Supplementary Text C for details). Specifically, we considered five chromosomes, each consisting of 200 genes of length 5,000 bp. The recombination rate is assumed to be 10^*−*7^ per site within a gene while the rate between adjacent genes is 0.005. Within each chromosome, three genes contained a selected site located at the center of the gene. Under this genomic architecture, sample data were simulated using SLiM v4.3 (Haller and Messer 2023).

In this situation, many neutral SNPs within genes harboring selected sites are expected to show strong genotype-environment associations due to linkage. As a consequence, SNP-level analyses naturally result in many false positives in a sense that many neutral loci are detected as significant. Therefore, the performance evaluation at the SNP-level is not informative in this setting, and we instead focused on gene-level analyses. For each gene, we used the most significant association among SNPs within the gene as a representative statistic and examined whether the resulting gene-level rank order successfully identifies adaptive genes using AUPRC. Since it is not straightforward to calculate gene-level *p*-values from SNP-level *p*-values (but see Yeaman *et al*. 2016; Booker *et al*. 2024), we here focus on the rank order and refrain from analyzing TPR and FDR.

We observe the largely consistent pattern with the linkage equilibrium simulations (Figure S6). The simple rank correlation test and SimGEA again showed high AUPRC, whereas LFMM2 and BayPass tended to show reduced performance under several parameter settings. This result suggests that linkage does not substantially change the relative performance of SimGEA.

### Performance under complex demography

Finally, we relaxed the assumption of simple demography by analyzing simulation data from Lotterhos and Whitlock (2015), deposited at https://doi.org/10.5061/dryad.mh67v. These simulations assume four demographic scenarios: isolation by distance, expansion from a single refugium, expansion from two refugia, and an island model. Each dataset consists of 9,900 neutral sites and approximately 100 selected sites, resulting in a higher proportion of adaptive SNPs than in our simulated datasets.

We compared the performance of SimGEA and other methods (rank correlation, LFMM2, and BayPass) using the three metrics described above (AUPRC, TPR, and FDR; Figure 5). We first focused on AUPRC to evaluate the ability of each method to rank truly adaptive sites near the top. Consistent with our previous results, SimGEA exhibited a pattern closely matching that of the simple rank correlation. LFMM2 showed comparable AUPRC values in most scenarios, but exhibited reduced AUPRC in some isolation-by-distance models. BayPass performed comparably to SimGEA under island models, but showed lower AUPRC under other demographic scenarios.

**Figure 5.**
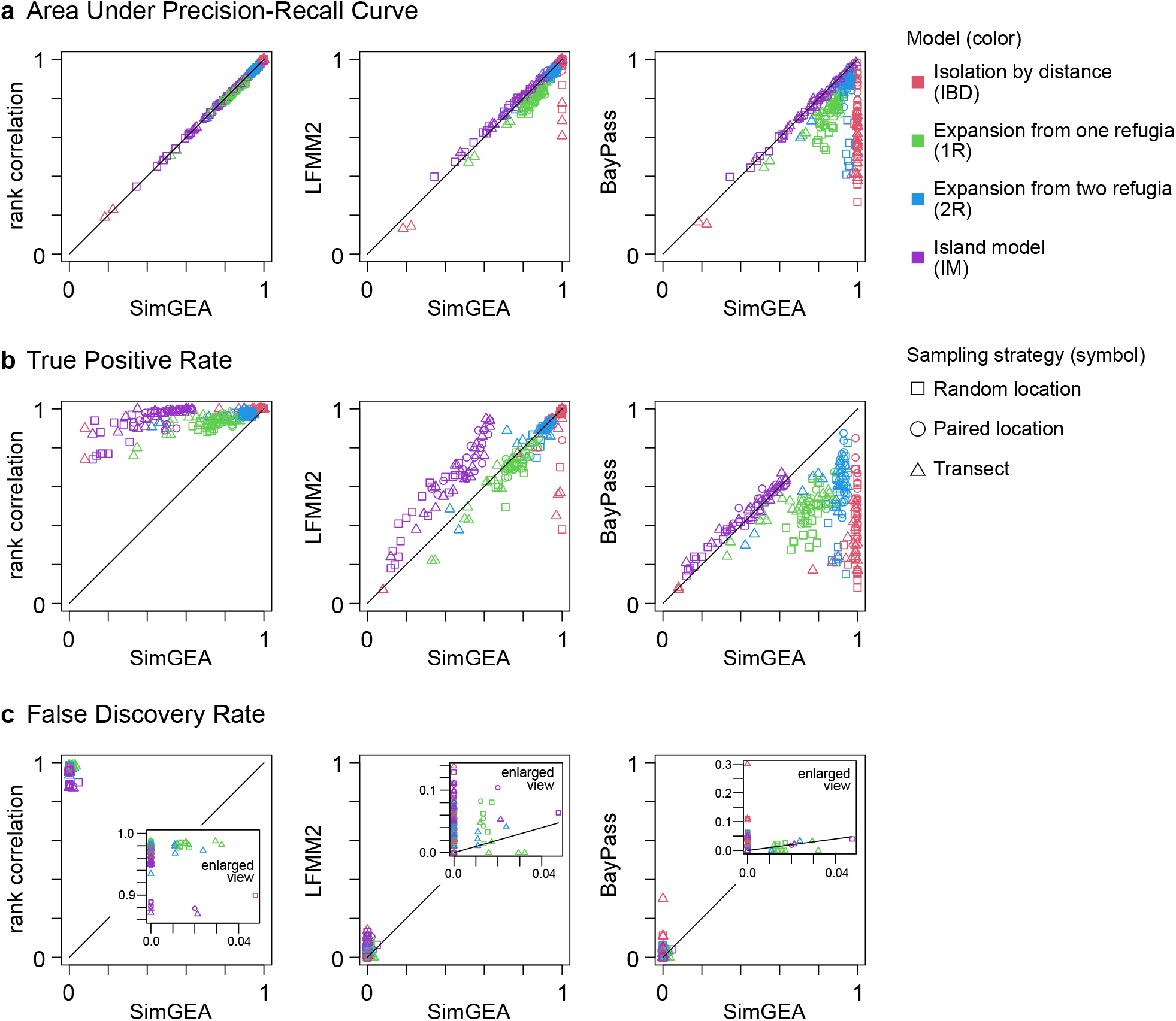
Performance under the different demographic models. The performance of a simple rank correlation, LFMM2, and Bay-Pass is compared against SimGEA using AUPRC (a), TPR (b), and FDR (c) . Simulation data from Lotterhos and Whitlock (2015) was used for this evaluation. Each color represents the different evolutionary scenarios while symbols show sampling strategies. Note that points with the same color and symbol do not mean that their underlying parameter is identical as they may differ in unrepresented parameters including the number of sampling locations and the number of sampled individuals per location.

We next calculated TPR and FDR using the same significance criteria as above. The simple rank correlation detected a large fraction of truly adaptive sites, but also exhibited elevated FDR due to its permissive significance threshold. LFMM2 showed higher power than SimGEA in island models, accompanied by a moderate increase in FDR. BayPass exhibited lower power outside island models and generally showed higher FDR than SimGEA. SimGEA consistently showed substantially lower FDR than our control target (0.05) and appeared overly conservative for this dataset. This behavior is partly due to the higher proportion of adaptive SNPs in this dataset (approximately 1%), which causes SimGEA to partially absorb adaptive signals into the inferred population structure. Note that this bias leads to more conservative estimates; thus loci detected by SimGEA are likely to be true positives.

## Discussion

Developing GEA methods that account for population structure is essential for reliably distinguishing adaptive signals from neutral evolution. Without correcting for population structure, GEA results in many false positives driven by isolation by distance (Meirmans 2012; De Mita *et al*. 2013). To address this issue, several methods have been developed (Frichot *et al*. 2013; Caye *et al*. 2019; Coop *et al*. 2010; Günther and Coop 2013; Gautier 2015; Forester *et al*. 2018); however, they often suffer from reduced statistical power and elevated false positives, likely due to collinearity between population structure and environmental variation (Yeaman *et al*. 2016; Capblancq *et al*. 2023).

Our new method, SimGEA, addresses this limitation by combining the estimation of migration parameters and a neutral simulation. By comparing observed associations with a null distribution generated from simulated neutral alleles, this framework provides *p*-values that effectively account for population structure. Validation using extensive spatial simulations shows that SimGEA applied to a simple rank correlation test achieves good statistical power while effectively controlling the false discovery rate. In contrast, existing methods with structure correction (i.e., LFMM2 and BayPass) tend to exhibit reduced power or inflated false discovery rates, consistent with previous studies. The performance difference is relatively large when the environmental map is partially gradient (Figure 4), which may often be the case in natural environments. These results suggest that SimGEA provides a reliable way to detect loci underlying local adaptation while minimizing the effect of population structure.

A key technical improvement enabling this approach is the fast estimation of a migration matrix for a large number of subpopulations. Although the migration rate can be estimated straightforwardly in simple Wright island models from *F*_*ST*_ *≈* 1/(4*Nm* + 1) (Wright 1931), this approximation breaks down in real-world metapopulations with interconnected and heterogeneous migration networks (Whitlock and McCauley 1999). Demographic inference methods have been widely used to estimate migration rates (Beerli and Felsenstein 2001; Beerli 2006; Gutenkunst *et al*. 2009; Excoffier and Foll 2011; Excoffier *et al*. 2021; Jouganous *et al*. 2017), but these methods are computationally demanding and limited to small numbers of subpopulations. Important exceptions are EEMS (Estimated Effective Migration Surfaces) (Petkova *et al*. 2016) and its subsequent extensions (Al-Asadi *et al*. 2019; Marcus *et al*. 2021; Shastry *et al*. 2025; Shen and Novembre 2026). These methods primarily aim to infer and visualize spatial patterns of migration rather than to construct a generative model that accurately reproduces allele-frequency covariance among sampled locations. Their suitability as baseline models for calibrating genotype-environment association tests is therefore not guaranteed. In contrast, our framework estimates population-scaled migration rates for ≥ 100 subpopulations that reproduce the empirical covariance pattern while maintaining computational feasibility. Although this method does not necessarily identify the true past demography as the parameter space in exploration is limited, it provides a useful null model that accounts for the neutral population structure reflected in the data.

Genome scans for locally adaptive variants provide insight into the genetic architecture underlying adaptation, because different genetic architectures are likely to produce different extent of repeatability in evolutionary patterns (Láruson *et al*. 2020). When the same genes are repeatedly involved in adaptation to a particular environment across different lineages, this suggests that the genetic routes to adaptation are limited, with little redundancy in the underlying genetic basis. In contrast, when adaptive variants show little overlap among independent lineages, this indicates that many different genotypes can confer adaptation to the focal environment, potentially reflecting high redundancy in the genetic basis of adaptation. The overlap among adaptive variants across different lineages can be formally tested using genome-scan results as input (*PicMin*; Booker *et al*. 2023), and this framework has been applied in many studies (Whiting *et al*. 2024; Nocchi *et al*. 2024; Soudi *et al*. 2023). In this approach, the power and accuracy of the genome scans is crucial, as the inclusion of false positives and the exclusion of true positives can underestimate repeatability (Booker *et al*. 2023). Our improvement in the GEA analysis may therefore contribute to accurate evaluation of repeated local adaptation within this framework.

Currently, many repeatability analyses rely on ranks of significance in GEA statistics rather than on absolute *p*-values (Booker *et al*. 2023), partly because absolute *p*-values can be poorly calibrated under complex demographic histories. However, because rank-based statistics do not quantify the absolute statistical significance of observed associations, well-calibrated *p*-values can enable more informative and potentially more powerful downstream analyses. Our new method provides structure-corrected *p*-values and could provide a basis for future extensions toward more powerful repeatability analyses in local adaptation studies.

In this study, we tested the performance of SimGEA mainly using two models: a two-dimensional stepping-stone model at equilibrium and the simulation datasets from Lotterhos and Whitlock (2015), which considered several non-equilibrium demographic scenarios. Although SimGEA showed consistently good performance compared with conventional methods, its overall performance should be carefully evaluated across a wider range of demographic scenarios. In natural populations, species distributions and environmental conditions change through time, as exemplified by repeated glacial cycles. Under such conditions, current genotype-environment relationships may not fully reflect the history of local adaptation, because allele-frequency clines shaped by adaptation to past environments may no longer show strong associations with contemporary environmental gradients. As different methods may show different power and robustness under such non-equilibrium dynamics, evaluating the performance of each method under diverse demographic and environmental histories will be important for interpreting GEA results from empirical data.

We here applied our simulation-based method to GEA analysis to account for the effects of population structure. However, because the core idea of our method is to compare the observed distribution of test statistics with a null distribution generated by neutral simulations, its applicability may extend beyond GEA analysis. Given that accounting for spatial structure is a fundamental challenge in a wide range of genomic analyses, extending this framework to other statistical tests represents a promising direction for future research.

## Data Availability

The code used to generate the simulation datasets (the island models and two-dimensional stepping-stone models) and its analysis is available at https://github.com/TSakamoto-evo/SimGEA/tree/main/simulation_analysis. The simulation dataset from Lotterhos and Whitlock (2015) analyzed in this study was downloaded from https://doi.org/10.5061/dryad.mh67v. The code used to run the full GEA analysis workflow in this study (SimGEA) is available at https://github.com/TSakamoto-evo/SimGEA/tree/main/simGEA_v1.1.

