## Supplementary materials for "A simulation-based method for genotype-environment association analysis"

August 19, 2026

#### Text A: Estimation of covariance matrix

In this section, we describe the detailed procedures and underlying derivations used to estimate a covariance matrix of allele frequencies (i.e., Step 1).

First, we modeled the sampling process using a binomial distribution. Let $p_{ij}$ and $b_{ij}$ be the allele frequency and the observed allele count at locus $i$ in subpopulation $j$, respectively. We denote by $a_j$ the number of sampled haplotypes in subpopulation $j$. As $b_{ij} \sim \mathrm{Binom}(a_j, p_{ij})$, the following expressions are obtained:

$$\begin{aligned}
\mathrm{E}[b_{ij}|p_{ij}] &= a_j p_{ij} \\
\mathrm{Var}[b_{ij}|p_{ij}] &= a_j p_{ij}(1-p_{ij}) \\
\mathrm{E}[b_{ij}b_{ik}|p_{ij},p_{ik}] &= a_j a_k p_{ij} p_{ik} \qquad (j \neq k).
\end{aligned} \tag{A1}$$

Using Equation 1 in the main text, unconditional mean and variance are given by

$$\begin{aligned}
\mathrm{E}[b_{ij}] &= a_j \bar{p}_i \\
\mathrm{Var}[b_{ij}] &= \mathrm{E}[\mathrm{Var}[b_{ij}|p_{ij}]] + \mathrm{Var}[\mathrm{E}[b_{ij}|p_{ij}]] \\
&= a_j \bar{p}_i(1-\bar{p}_i)(1-\Lambda_{jj}) + a_j^2 \bar{p}_i(1-\bar{p}_i)\Lambda_{jj} \\
\mathrm{E}[b_{ij}b_{ik}] &= a_j a_k \mathrm{E}[p_{ij}p_{ik}] \\
&= a_j a_k \left[\bar{p}_i(1-\bar{p}_i)\Lambda_{jk} + \bar{p}_i^2\right] \qquad (j \neq k) \\
\mathrm{Cov}[b_{ij}, b_{ik}] &= \mathrm{E}[b_{ij}b_{ik}] - \mathrm{E}[b_{ij}]\mathrm{E}[b_{ik}] \\
&= a_j a_k \bar{p}_i(1-\bar{p}_i)\Lambda_{jk} \qquad (j \neq k)
\end{aligned} \tag{A2}$$

Let $z_{ij} = b_{ij}/a_j$ be the observed allele frequency in the sample. Using the above equations, we obtain

$$\begin{aligned}
\mathrm{E}[z_{ij}] &= \bar{p}_i \\
\mathrm{Var}[z_{ij}] &= \frac{\bar{p}_i(1-\bar{p}_i)(1-\Lambda_{jj})}{a_j} + \bar{p}_i(1-\bar{p}_i)\Lambda_{jj} \\
\mathrm{Cov}[z_{ij}, z_{ik}] &= \bar{p}_i(1-\bar{p}_i)\Lambda_{jk} \qquad (j \neq k)
\end{aligned} \tag{A3}$$

We define the mean allele frequency in the sample across subpopulations as $\bar{z}_i = \frac{\sum_j z_{ij}}{n}$ (rather than $\frac{\sum_j b_{ij}}{\sum_j a_j}$). Noting that each row and column of $\Lambda$ sums to zero by definition, we obtain

$$\begin{aligned}
\mathrm{E}[\bar{z}_i] &= \bar{p}_i \\
\mathrm{Var}[\bar{z}_i] &= \frac{1}{n^2}\left[\sum_j \mathrm{Var}[z_{ij}] + \sum_{j \neq k} \mathrm{Cov}[z_{ij}, z_{ik}]\right] \\
&= \frac{1}{n^2}\sum_j \frac{\bar{p}_i(1-\bar{p}_i)(1-\Lambda_{jj})}{a_j}.
\end{aligned} \tag{A4}$$

From these, Equation 2 is derived:

$$
\begin{aligned}
\mathrm{E}[z_{ij} - \bar{z}_i] &= 0 \\
\mathrm{Var}[z_{ij} - \bar{z}_i] &= \mathrm{Var}[z_{ij}] - \frac{2}{n}\sum_k \mathrm{Cov}[z_{ij}, z_{ik}] + \mathrm{Var}[\bar{z}_i] \\
&= \left(1 - \frac{2}{n}\right)\frac{\bar{p}_i(1-\bar{p}_i)(1-\Lambda_{jj})}{a_j} + \bar{p}_i(1-\bar{p}_i)\Lambda_{jj} + \frac{1}{n^2}\sum_{l=1}^{n}\frac{\bar{p}_i(1-\bar{p}_i)(1-\Lambda_{ll})}{a_l} \\
\mathrm{Cov}[z_{ij} - \bar{z}_i, z_{ik} - \bar{z}_i] &= \mathrm{Cov}[z_{ij}, z_{ik}] - \frac{1}{n}\sum_l \mathrm{Cov}[z_{ij}, z_{il}] - \frac{1}{n}\sum_l \mathrm{Cov}[z_{ik}, z_{il}] + \mathrm{Var}[\bar{z}_i] \\
&= \bar{p}_i(1-\bar{p}_i)\Lambda_{jk} - \frac{1}{n}\frac{\bar{p}_i(1-\bar{p}_i)(1-\Lambda_{jj})}{a_j} - \frac{1}{n}\frac{\bar{p}_i(1-\bar{p}_i)(1-\Lambda_{kk})}{a_k} + \frac{1}{n^2}\sum_{l=1}^{n}\frac{\bar{p}_i(1-\bar{p}_i)(1-\Lambda_{ll})}{a_l} \qquad (j \neq k).
\end{aligned}
\tag{A5}
$$

Equation 3 is then derived by assuming $\bar{p}_i \approx \bar{z}_i$. By substituting $S_{jk}$ values obtained from the sample (Equation 4), Equation 3 represents a system of equations for $\Lambda_{jk}$. The first line of Equation 3 gives $n$ equations for $\Lambda_{jj}$, which can be numerically solved. These values are then substituted into the second line to calculate $\Lambda_{jk}$ for $j \neq k$.

Finally, since true $\mathbf{\Lambda}$ must be a positive semi-definite matrix, we modified the estimated $\mathbf{\Lambda}_{\mathrm{obs}}$ when it is not positive semi-definite by replacing negative eigenvalues with zero, assuming that this deviation is due to the sampling noise.

### Text B: Inference of migration matrix

In this section, we describe the detailed procedure for estimating the migration matrix (i.e., Step 2). Our aim is to infer a migration matrix that reproduces the estimated covariance matrix of allele frequencies ($\mathbf{\Lambda}_{\mathrm{obs}}$), while restricting the search space to an island model with equal haploid population size $N$ and symmetric migration rates, $m_{ij} = m_{ji}$.

#### Calculation of $\mathbf{\Lambda}(\boldsymbol{M})$

Let $\boldsymbol{M}$ be an $n \times n$ matrix such that $M_{ij} = 2Nm_{ij}$ for $i \neq j$, and $M_{ii} = -\sum_{j \neq i} M_{ij}$. The equilibrium state of the focal island model is fully determined by $\boldsymbol{M}$ (Wakeley, 2009).

To search for an optimal migration matrix $\boldsymbol{M}$, we first need to know the relationship between $\boldsymbol{M}$ and the resulting covariance matrix $\mathbf{\Lambda}$. For this purpose, we employ structured coalescent theory (Wakeley, 2009). Let $T_{ij}$ be the expected coalescent time between a sample from subpopulation $i$ and a sample from subpopulation $j$, measured in units of the local population size $N$. The expected coalescent times satisfy the following system of equations (Wakeley, 2009):

$$
\begin{aligned}
1 &= (1 - M_{ii})T_{ii} - \sum_{j \neq i} M_{ij}T_{ij} \\
2 &= -(M_{ii} + M_{jj})T_{ij} - \sum_{k \neq i} M_{ik}T_{jk} - \sum_{k \neq j} M_{jk}T_{ik}
\end{aligned}
\tag{B1}
$$

Let $U = 2Nu$ be the population-scaled per-site mutation rate. The expected heterozygosity between two samples drawn from subpopulations $i$ and $j$ is given by $H_{ij} = UT_{ij}$. We also denote the allele frequency in subpopulation $i$ by $x_i$, and its global mean by $\bar{x} = 1/n \times \sum_i x_i$. Using these notations, we obtain

$$\begin{aligned}
H_{ij} &= \mathrm{E}[x_i(1-x_j) + (1-x_i)x_j] \\
&= \mathrm{E}[x_i] + \mathrm{E}[x_j] - 2\mathrm{E}[x_i x_j] \\
\mathrm{E}[(x_i-\bar{x})(x_j-\bar{x})] &= \mathrm{E}[x_i x_j] - \frac{1}{n}\sum_k \mathrm{E}[x_i x_k] - \frac{1}{n}\sum_k \mathrm{E}[x_j x_k] + \frac{1}{n^2}\sum_{k,l}\mathrm{E}[x_k x_l] \\
&= -\frac{1}{2}\left(H_{ij} - \frac{1}{n}\sum_k H_{ik} - \frac{1}{n}\sum_k H_{jk} + \frac{1}{n^2}\sum_{k,l} H_{kl}\right) \\
\mathrm{E}[\bar{x}(1-\bar{x})] &= \frac{1}{2n^2}\sum_{k,l} H_{kl}
\end{aligned} \tag{B2}$$

Based on Equation 1 in the main text, $\Lambda_{ij}$ is then assumed to follow

$$\begin{aligned}
\Lambda_{ij} &= \frac{\mathrm{E}[(x_i-\bar{x})(x_j-\bar{x})]}{\mathrm{E}[\bar{x}(1-\bar{x})]} \\
&= -\frac{n^2 T_{ij} - n\sum_k T_{ik} - n\sum_k T_{jk} + \sum_{k,l} T_{kl}}{\sum_{k,l} T_{kl}}
\end{aligned} \tag{B3}$$

Equation B3 shows that the expected covariance matrix $\boldsymbol{\Lambda}$ can be obtained once the expected pairwise coalescent times are determined by Equation B1. Since Equation B1 constitutes a system of $n(n+1)/2$ linear equations, $T_{ij}$ can numerically be determined although solving it becomes computationally expensive for large $n$. In our implementation, we compute the solution efficiently using a sparse matrix solver from the Eigen C++ library.

#### Optimization

In the optimization step, we search for a migration matrix $\boldsymbol{M}$ that minimizes the distance between the theoretical covariance matrix $\boldsymbol{\Lambda}_{\mathrm{theor}}(\boldsymbol{M})$ and the observed covariance matrix $\boldsymbol{\Lambda}_{\mathrm{obs}}$. The distance between two matrices $\boldsymbol{X}$ and $\boldsymbol{Y}$ is measured as

$$D(\boldsymbol{X},\boldsymbol{Y}) = \sum_{i<j}(x_{ij} - y_{ij})^2.$$

We parameterize the migration rates as $M_{ij} = \log(1+\exp(\mu_{ij}))$ to ensure $M_{ij} > 0$, and optimize the $n(n-1)/2$ parameters $\mu_{ij}$ using the AdamW algorithm (Loshchilov and Hutter, 2017). Due to the high computational cost of evaluating $\boldsymbol{\Lambda}(\boldsymbol{M})$, it is crucial to reduce the number of the calculations of $\boldsymbol{\Lambda}(\boldsymbol{M})$ during optimization. To this end, we employ the pseudo-gradient approach described in the main text.

The optimization procedure at each step is summarized as follows:

- Obtain $\boldsymbol{M}$ from the current values of $\mu_{ij}$.
- Compute $\boldsymbol{\Lambda}(\boldsymbol{M})$ using Equations B1 and B3.
- Evaluate the distance $D(\boldsymbol{\Lambda}(\boldsymbol{M}), \boldsymbol{\Lambda}_{\mathrm{obs}})$.
- Compute the pseudo-gradient as

$$\begin{aligned}
\text{pseudo-grad}_{ij} \equiv \frac{\partial D}{\partial \mu_{ij}} &\approx \frac{dM_{ij}}{d\mu_{ij}} \sum_{k<l}\left(\Lambda_{\mathrm{theor},kl} - \Lambda_{\mathrm{obs},kl}\right)\Delta\Lambda^{(ij)}_{\mathrm{theor},kl} \\
&= \frac{1}{1+\exp(-\mu_{ij})}\sum_{k<l}\left(\Lambda_{\mathrm{theor},kl} - \Lambda_{\mathrm{obs},kl}\right)\Delta\Lambda^{(ij)}_{\mathrm{theor},kl}.
\end{aligned} \tag{B4}$$

See also Equation 5 in the main text.

- Update $\mu_{ij}$ using the AdamW algorithm with pseudo-grad$_{ij}$.

For the AdamW optimizer, we use the following parameter values throughout this study: $\beta_1 = 0.9$, $\beta_2 = 0.999$, learning rate $\alpha = 0.001$, weight decay $\lambda = 0.01$, and $\epsilon = 10^{-8}$.

Let $D(t)$ denote the value of $D(\boldsymbol{\Lambda}(\boldsymbol{M}), \boldsymbol{\Lambda}_{\text{obs}})$ at optimization step $t$. The optimization is terminated when any of the following conditions is satisfied:

- After at least 1000 steps, ten successive steps satisfy $\frac{2D(t-1)-2D(t)}{2D(t-1)+\tau_1} < \tau_1$ and $2D(t-1) - 2D(t) < \frac{2D(0)-2D(1000)}{1000}\tau_2$.
- At $t = 1000k$ ($k \geq 2$, $k \in \mathbb{N}$), $\frac{2D(t-1000)-2D(t)}{2D(t-1000)+\tau_1} < \tau_1$.
- The total number of optimization steps reaches $\tau_3$.

In this study, we set $\tau_1 = 10^{-6}$, $\tau_2 = 0.01$, and $\tau_3 = 200{,}000$.

The estimated migration matrix is then used to perform coalescent simulations. Neutral alleles generated by these simulations are expected to exhibit population structure similar to that of the sampled data, and therefore provide a baseline for detecting alleles under selection.

### Text C: Details of our simulated dataset

#### Island model

We used simulated datasets to evaluate the ability of our method to recover the true population structure. For this purpose, we first considered an island model with $n$ subpopulations of equal population size $N$ and symmetric migration rates $m_{ij}$. In this setting, the true model is within the search space of our optimization step, thus providing an opportunity to evaluate our method under an ideal situation.

Firstly, a random migration matrix is generated, where the overall strength of migration is controlled by a parameter $M_{\text{tot}}$. For each pair $(i, j)$ with $i > j$, the migration parameter $M_{ij} = 2Nm_{ij}$ was independently drawn from an exponential distribution with mean $M_{\text{tot}}/n$. This matrix was then used to simulate neutral alleles using a coalescent simulator, in which $a$ haploid samples (or equivalently $a/2$ diploid samples) were taken from each subpopulation across $S$ segregating loci. In our simulations, we set $a = 8$ and $S = 10^5$. To reduce the linkage among variants due to the multiple mutations on a tree, we set the mutation rate so that 0.25 mutations, on average, arise from a single tree. We then keep sampling coalescent trees until the total number of mutations reach $S$.

We considered three values of $n$ and $M_{\text{tot}}$, namely $n = 10, 30$, and 100, and $M_{\text{tot}} = 1, 10$, and 100. For each of the nine parameter combinations, we generated 20 independent migration matrices to assess how accurately the estimated population structure recovers the true underlying structure.

#### 2D stepping-stone model under linkage equilibrium

We next consider more realistic spatial models and evaluate the ability to detect adaptive variants in comparison with previous methods. To this end, we consider a two-dimensional stepping stone model with $14 \times 14$ subpopulations with diploid subpopulation size $N = 100$. To start from a simple situation, linkage equilibrium is assumed. Migration occurs between adjacent subpopulations (von-Neumann neighborhood) at the rate of $m$ per direction per generations. In this study, we consider two migration rates: $m = 0.04$ (high) and $m = 0.01$ (low).

We consider both neutral and selected variants. New neutral variants arise at the rate of $\mu_n = 0.05$ per genome per generation under the infinite-site model. Allele frequencies at polymorphic sites are simulated following Wright–Fisher model: by letting the expected allele frequency at the coordinate $(x, y)$ at generation $t$ be $p'_{xy}(t)$, the allele frequency at generation $t$ is determined by $p_{xy}(t) \sim \text{Binom}(2N, p'_{xy}(t))$. For example, the allele frequency at the subpopulation with coordinate $(5, 5)$ is calculated as

$$\begin{aligned} &p'_{5,5}(t+1) = (1-4m)p_{5,5}(t) + mp_{4,5}(t) + mp_{6,5}(t) + mp_{5,4}(t) + mp_{5,6}(t) \\ &p_{5,5}(t+1) \sim \text{Binom}(2N, p'_{5,5}(t+1)), \end{aligned} \tag{C1}$$

and that at the one with (0, 0) (i.e., at the corder of the habitat) is calculated as

$$\begin{aligned} p'_{0,0}(t+1) &= (1-2m)p_{0,0}(t) + mp_{1,0}(t) + mp_{0,1}(t) \\ p_{0,0}(t+1) &\sim \text{Binom}(2N, p'_{0,0}(t+1)). \end{aligned} \tag{C2}$$

We also consider 15 selected loci, consisting of five loci each with strong ($s_E = 0.01$), intermediate ($s_E = 0.003$), and weak selection ($s_E = 0.001$). These loci are subject to environment dependent selection, where selection coefficient at the coordinate $(x, y)$ is given by the product of $s_E$ and environment $E(x, y)$. In this study, we considered three environmental maps with different degree of spatial autocorrelation (Figure 4a), where all maps are centralized (mean 0) and normalized (variance 1) across the habitat. The mutation rate for these loci is assumed to be $\mu_s = 10^{-7}$ per locus per generation. Allele frequency is then calculated following the equation:

$$\begin{aligned} p'_{xy}(t+1) &= p_{xy}(t) + \underbrace{s_E E(x,y)p_{xy}(t)(1-p_{xy}(t))}_{\text{selection}} + \underbrace{\mu_s(1-2p_{xy}(t))}_{\text{mutation}} + \underbrace{\sum_{(i,j)\in \text{adj}(x,y)} m(p_{ij}(t)-p_{xy}(t))}_{\text{migration}} \\ p_{xy}(t+1) &\sim \text{Binom}(2N, p'_{xy}(t+1)). \end{aligned} \tag{C3}$$

To ensure that the system enters the equilibrium state, we took the samples after 400,000 generations. For each simulation run, five different sampling schemes are applied (1 ind.$\times$160 loc., 2 ind.$\times$80 loc., 4 ind.$\times$40 loc., 8 ind.$\times$20 loc., and 16 ind. $\times$ 10 loc.). Sampling locations are randomly determined for each replication and sampling scheme, so they are not necessarily nested across the schemes. For each of six combinations of migration rates and environmental maps, 50 simulation replicates were conducted. We removed loci with the minor allele frequency MAF $< 0.05$ from the analysis. The observed patterns of $F_{ST}$ and the number of loci in the final dataset are summarized in Figure S4.

#### 2D stepping-stone model with linkage

We next consider a gene-based model to represent linkage disequilibrium. While the basic setting is the same as the linkage equilibrium model, the genomic positions are explicitly modeled (see Figure S7). We assumed five chromosomes in which 200 genes with length 5 kb exist. The recombination rate within a gene is $r_1 = 10^{-7}$ per site, while the recombination rate between two adjacent genes is $r_2 = 0.005$, resulting in the chromosome-wide recombination rate of $\approx 1$. In the middle of 33rd, 100th, and 167th genes, a selected site exists where mutation arises at the rate of $\mu_s = 10^{-7}$. Three selected loci have $s_E$ corresponding to strong (0.01), intermediate (0.003), and weak (0.001), respectively, but their positional order is determined randomly. The neutral mutation rate is assumed to be $10^{-8}$ per site per generation, resulting in $\mu_n = 0.05$ across the genome.

We implemented this model using SLiM v.4.3 (Haller and Messer, 2023). To achieve efficient computation, tree recoding mode was employed, and neutral mutations are added using *tskit* (Wong *et al.*, 2024). Although we run SLiM simulations for 400,000 generations, some genomic regions do not achieve a full coalescence. For these trees, ancestries are simulated until the full coalescence while assuming neutrality by using a recapitate function in pyslim. We finally removed loci with the minor allele frequency MAF $< 0.05$ from the analysis.

#### Simulation data from Lotterhos and Whitlock (2015)

To see the performance in non-equilibrium evolutionary dynamics, we used the simulation dataset from Lotterhos and Whitlock (2015). The data was downloaded from `https://doi.org/10.5061/dryad.mh67v`. In that study, the authors assumed various demographic histories, including expansion from refugia, as well as multiple sampling strategies to examine their effect on GEA performance. Using their LFMM-style genotype and environmental data, BayPass-style files are generated and used to run BayPass and SimGEA.

### Text D: GEA methods

#### LFMM2

To run LFMM2 (Caye *et al.*, 2019), we used the *LEA* package (v 3.19.8) in R (Frichot and François, 2015). For each dataset, we first inferred the number of latent factors by running the snmf function for values of $K$ ranging from 1 to 15. If a local minimum in cross-entropy was observed, the corresponding value of $K$ was used for LFMM2. When cross-entropy reached its minimum near the upper bound of the tested range (i.e., $K = 14$ or 15), we selected $K$ based on the elbow point of the cross-entropy curve. LFMM2 was then run using the selected number of latent factors, and $p$-values are used to define adaptive alleles while applying correction for multiple test.

#### BayPass

We run BayPass (v 2.4.1 Gautier, 2015) with the standard covariate model using an Importance Sampling approximation (i.e., default mode). Significance at each SNP is evaluated based on Bayes Factor. Each MCMC is run using default parameters.

#### Correlation test without correcting population structure

As an example of simple correlation analyses, we examined the rank-biserial correlation between environmental values and allele presence-absence (represented as 0 or 1) at each locus (Wendt, 1972). Because the rank-biserial correlation used here is mathematically derived from the Mann–Whitney U statistic, the corresponding $p$-values were obtained from Mann–Whitney U tests with correction for tied observations.

#### SimGEA

We used the SimGEA approach to calibrate $p$-values derived from the rank-biserial correlation. First, neutral alleles were simulated under a structured coalescent model parameterized by the estimated migration matrix. For each possible minor-allele count, $n_{\text{mut}}$ mutations were collected from numerous simulated coalescent trees. Each SNP in the empirical data was then compared with simulated neutral alleles with similar allele frequencies.

For this comparison, we employed an allele-count-based sliding window approach. Let $x$ denote the total number of sampled alleles per SNP site and $w$ the specified window width on the allele-count scale. Because the possible minor-allele counts range from 1 to $\lfloor x/2 \rfloor$, we considered $\left\lfloor \frac{x}{2} \right\rfloor - w + 1$ overlapping windows, as shown in Figure S8a. Each empirical SNP was matched to the window with the closest mean minor-allele count. For SNPs near the boundaries of the allele-count range, the window was shifted to retain the specified width.

For each window, we applied the rank-biserial correlation to simulated neutral datasets and obtained the distribution of $Z$ values. We focused on $Z$ values rather than $p$-values, because $Z$ values show a one-to-one correspondence with $p$-values while exhibiting a unimodal distribution. This distribution was then used as the null distribution after applying smoothing, as described below.

To obtain a smoothed null distribution, we applied kernel density estimation (KDE) with Gaussian kernels to the bulk of the distribution, defined as values up to the 99.5th percentile. For the tail of the distribution, KDE may perform poorly despite this region being critical for assessing statistical significance. We therefore modeled the tail using extreme value fitting based on the generalized Pareto distribution (Knijnenburg *et al.*, 2009). A fitted generalized Pareto distribution with a negative shape parameter has a finite upper endpoint, and an unconstrained fit can therefore assign zero or undefined tail probabilities to sufficiently large $Z$ values. This is undesirable in our application because truly adaptive variants may show substantially stronger associations (i.e., larger $Z$ values) than those observed among the simulated neutral variants. To avoid this problem, we constrained the fitted distribution. For each environmental variable and focal minor-allele count, we calculated the largest and second-largest possible $Z$-values, denoted by $Z_{\text{1st}}$ and $Z_{\text{2nd}}$, respectively. Because the possible $Z$-values are discrete, whereas the generalized Pareto distribution is continuous, we used the difference $\Delta_Z = Z_{\text{1st}} - Z_{\text{2nd}}$ to approximate the effective width of the largest Z-value in the continuous distribution. We then required the fitted distribution to remain defined up to at least $Z_{\text{1st}} + \Delta_Z$ (Figure S8b). As this procedure can make the fitted distribution heavier-tailed, this correction is expected to yield more conservative $p$-values. This procedure enables stable estimation of extreme $p$-values beyond the range directly represented by the simulated neutral variants, while ensuring that the fitted tail covers the possible range of the test statistic. In Supplementary Text E, we test whether this fitting results in the accurate $p$-values in general.

In the main analysis, the window size $w = 9$ was chosen so that each window spanned an allele-frequency range of 0.025, and $n_{\mathrm{mut}} = 11{,}112$ was set so that each window contains $\sim 10^5$ simulated neutral variants.

##### The method of Goel et al. (2026, bioRxiv)

Goel *et al.* (2026) recently proposed a genotype-environment association (GEA) method based on a hierarchical Bayesian model. To improve computational efficiency, they recommended a two-step analysis in which the migration matrix is first estimated and then treated as a known input in the downstream evaluation of genotype-environment associations. In both steps, inference is performed by Markov chain Monte Carlo (MCMC) implemented in Stan. The original framework can incorporate genotype uncertainty arising from low-coverage sequencing data, whereas our simulated datasets consist of error-free allele counts. We therefore implemented a simplified version of their migration-estimation component by omitting the genotype-inference component. In addition, their migration model allows covariates, such as geographic or landscape distances between subpopulations, to be incorporated when estimating migration rates.

For island-model simulations, we applied only the first step of this framework to obtain an estimated migration matrix. Because these simulations do not involve explicit geographic positions, no covariates were provided. We ran the MCMC using Stan v2.38.0 with three chains, 1000 warm-up iterations, and 1000 sampling iterations. Due to the high computational cost, we performed this analysis only for cases with $n = 10$ and 30 subpopulations.

We did not apply this approach to the stepping-stone simulations because of its long computational time. Thus, a comprehensive evaluation of the full framework of Goel *et al.* (2026) is left for future studies.

#### Text E: Validation of the $p$-value estimation method

In our analysis, we employed Gaussian kernel smoothing for the bulk and generalized Pareto fitting for the tail of the $Z$-value distribution to obtain well-calibrated $p$-values from a limited number of simulated variants. However, $p$-values for extreme $Z$ values may be less precise because their estimation requires extrapolation beyond the range directly represented by the simulated neutral alleles.

To examine the extent of this issue, we conducted the following validation. We first prepared a migration matrix and an environmental map. We next simulated $10^8$ variants for each possible allele count and calculated the empirical distribution of $Z$-values. We then applied our $p$-value estimation procedure to these $Z$-values (Supplementary Text D) and checked whether the empirical $p$-values were accurately recovered. In the p-value estimation, we used a frequency-window width of 0.025 and $10^5$ simulated variants per window, consistent with our main analysis. To evaluate variation among estimates, we repeated this p-value estimation 100 times.

Figure S9-S11 compares the estimated and empirical $p$-values for a subset of allele counts. Black solid lines represent median $p$-values, the gray regions indicate the 5th-95th percentile ranges, and the dashed lines represent the minimum and the maximum values across 100 replicates. The figure shows that our fitting generally works well, and that, as expected, deviations tended to produce conservative $p$-values. We observed similar overall trends for other allele counts. These results suggest that our strategy of estimating the $Z$-value distribution from a limited number of neutral alleles is useful for our purpose.

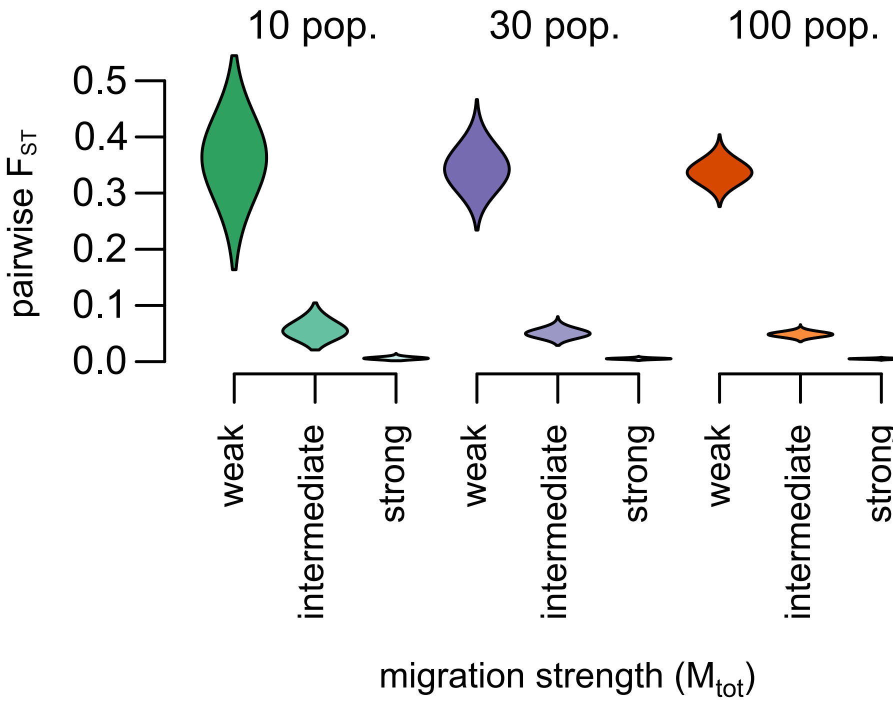


Figure S1: The distribution of $F_{ST}$ between pairs of subpopulations in each parameter setting. $F_{ST}$ is calculated as $F_{ST} = (H_T - H_W)/H_T$ (Nei, 1973; Slatkin, 1991).

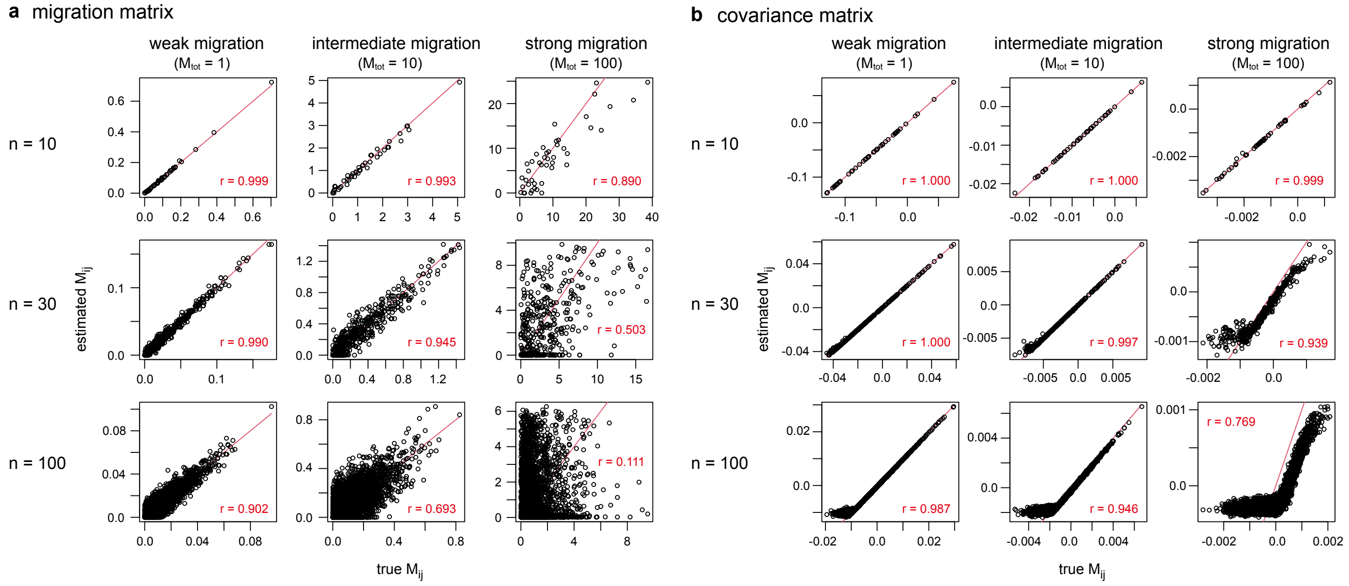


Figure S2: Comparison between the target and estimated matrices. (a) Comparison of $M_{ij}$ across parameter settings. For each panel, the result of one replicate is shown. (b) Covariance matrices are compared in the same run as the panels (a).

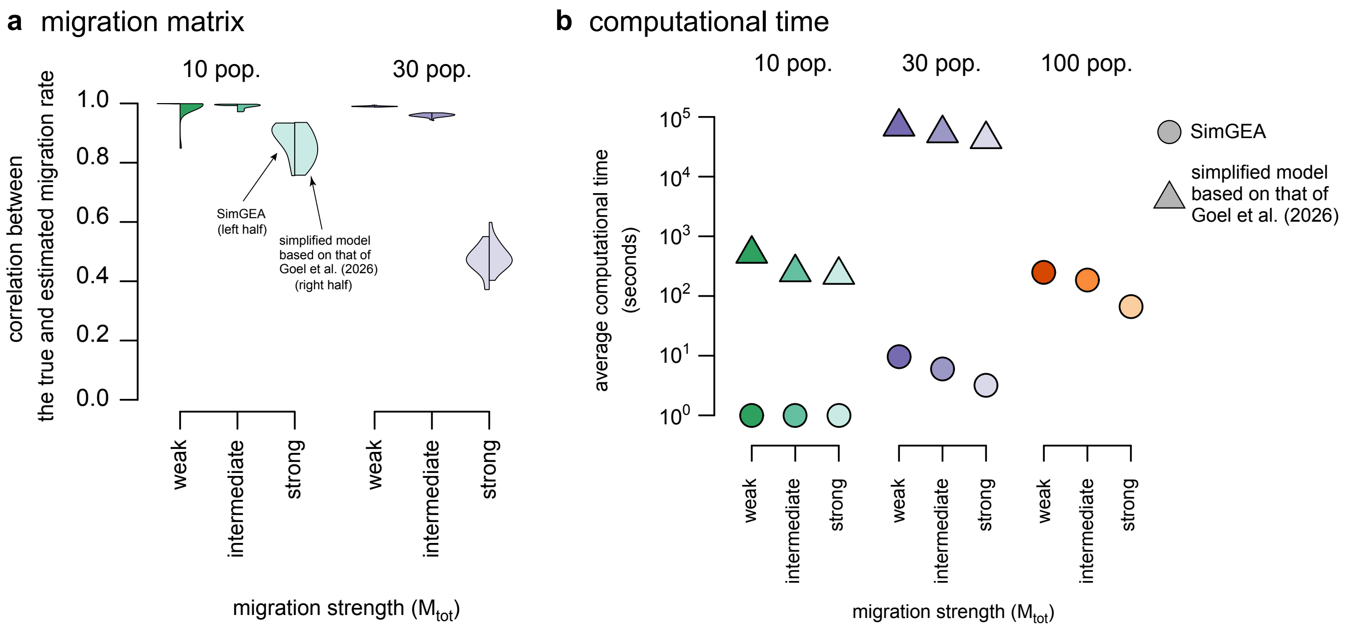


Figure S3: Comparison between SimGEA and the simplified MCMC model based on Goel *et al.* (2026). (a) Correlation coefficients in migration rates. In each violin plot, the left half shows the distribution produced by SimGEA across 20 replicates, while the right half represents the results of the MCMC model based on Goel *et al.* (2026). (b) For both methods, the programs were run on the computer cluster SHIROKANE at the Human Genome Center, The University of Tokyo, using a single thread per run. Since SHIROKANE is a shared computing environment, the cluster conditions and CPU types may vary to some extent across runs; therefore, the reported execution times should be regarded as rough indicators.

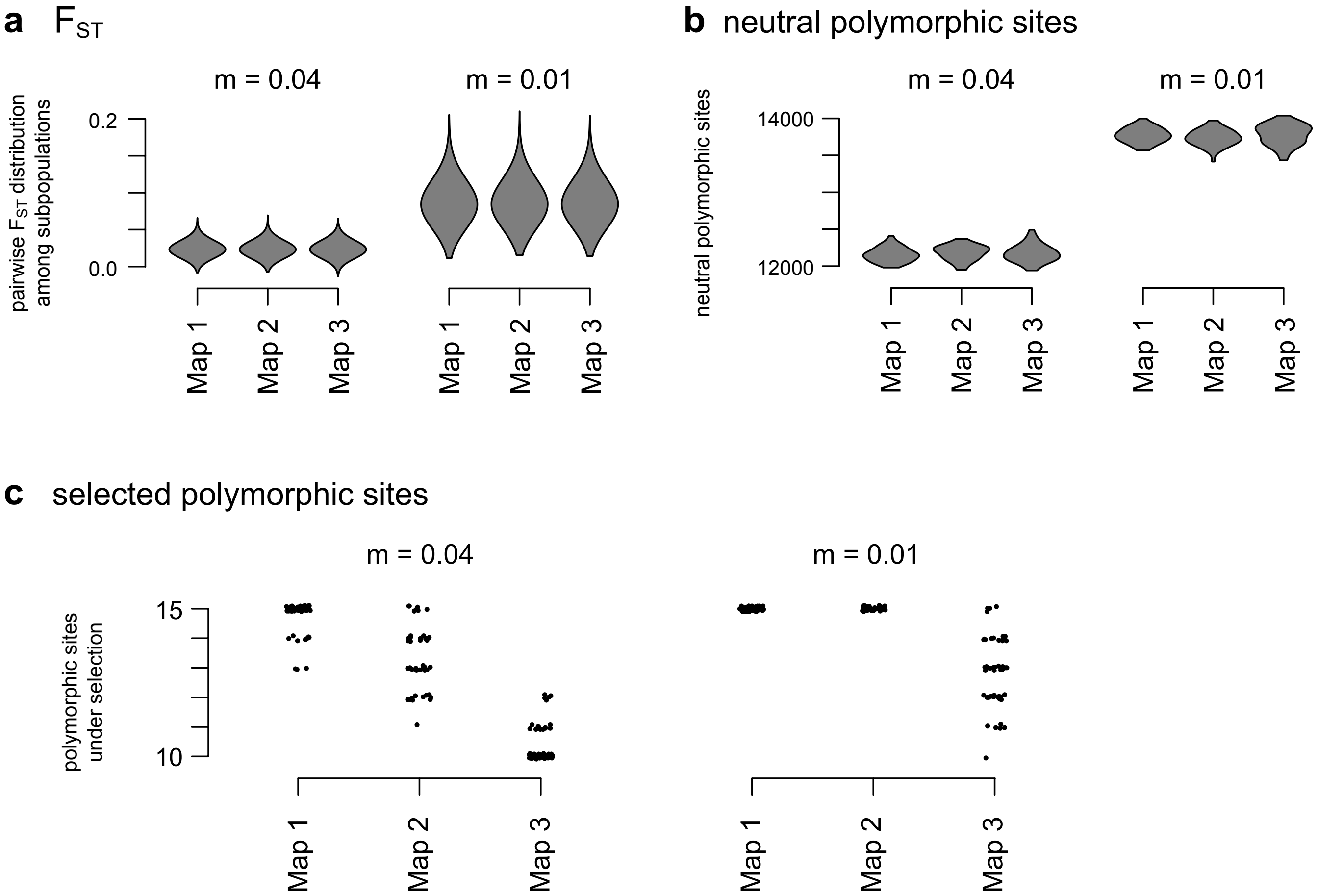


Figure S4: The distribution of pairwise $F_{ST}$ among subpopulations (a), the number of neutral polymorphic sites (b), and the number of selected polymorphic sites in our two-dimensional stepping stone model under linkage equilibrium. These statistics are calculated after applying a MAF filter at MAF $\geq 0.05$.

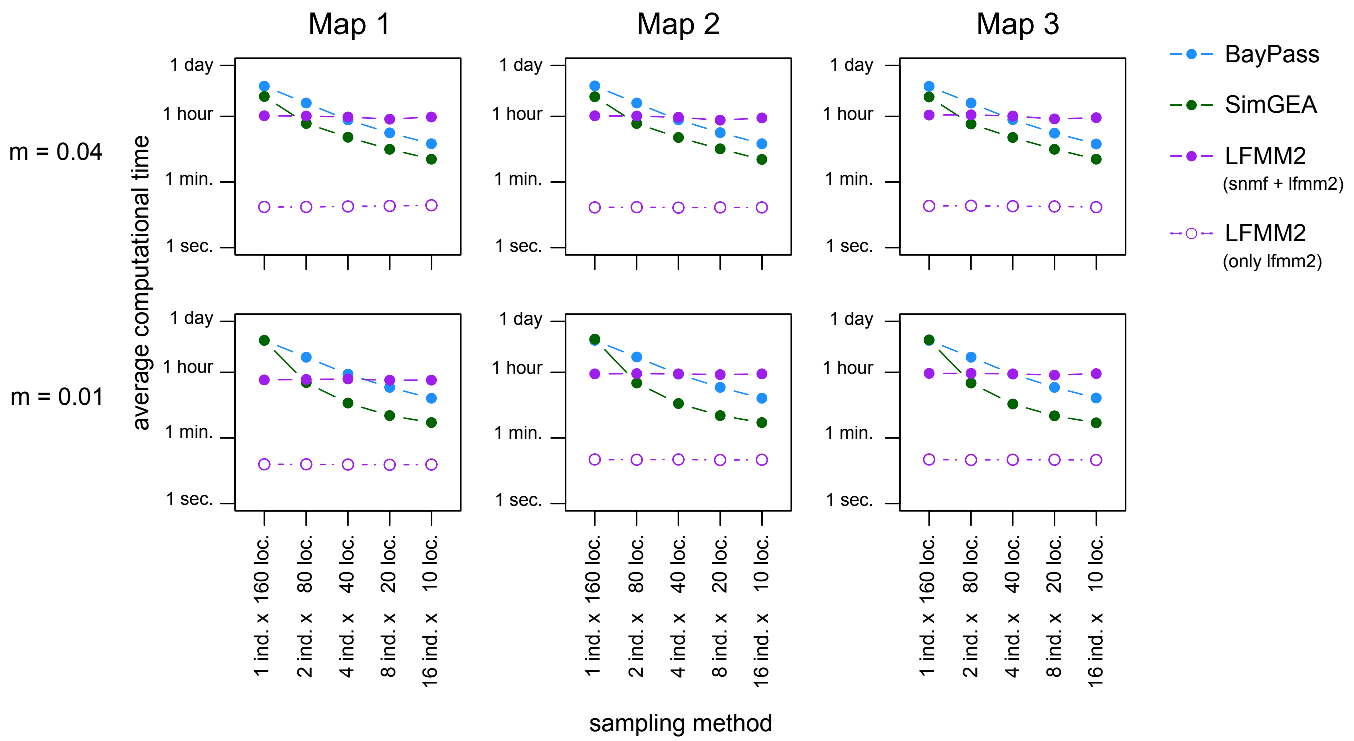


Figure S5: Comparison of computational time among BayPass, SimGEA, and LFMM2. For LFMM2, the estimation of the number of latent factors ($K$) using snmf is the time-consuming process in our analysis, so we provide both the total time and the time for running lfmm2 function after specifying $K$.

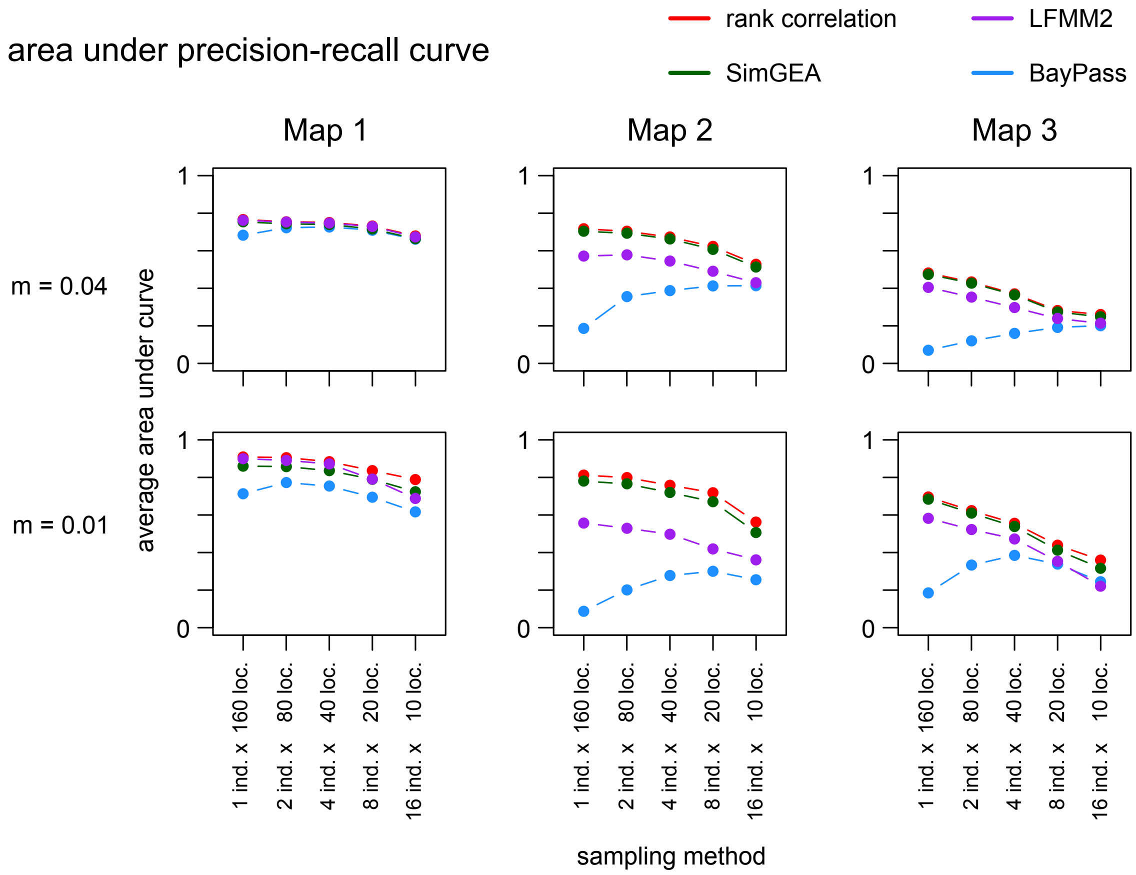


Figure S6: Performance comparison in the presence of linkage disequilibrium.

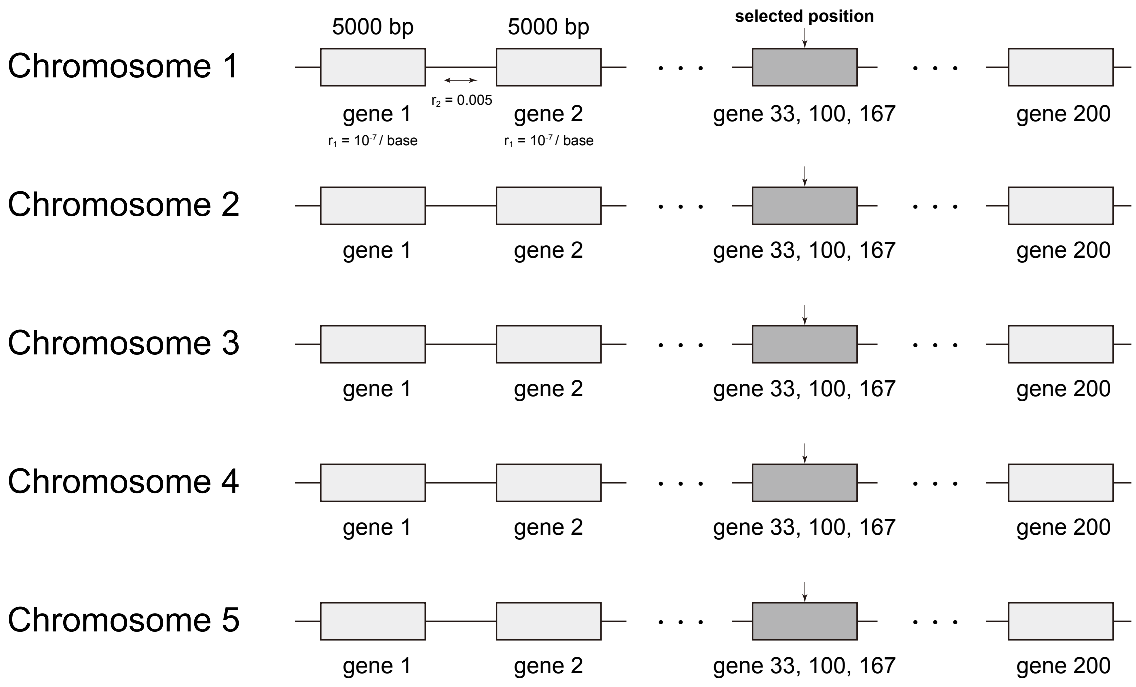


Figure S7: Schematic view of the modeled genome in simulations with linkage.

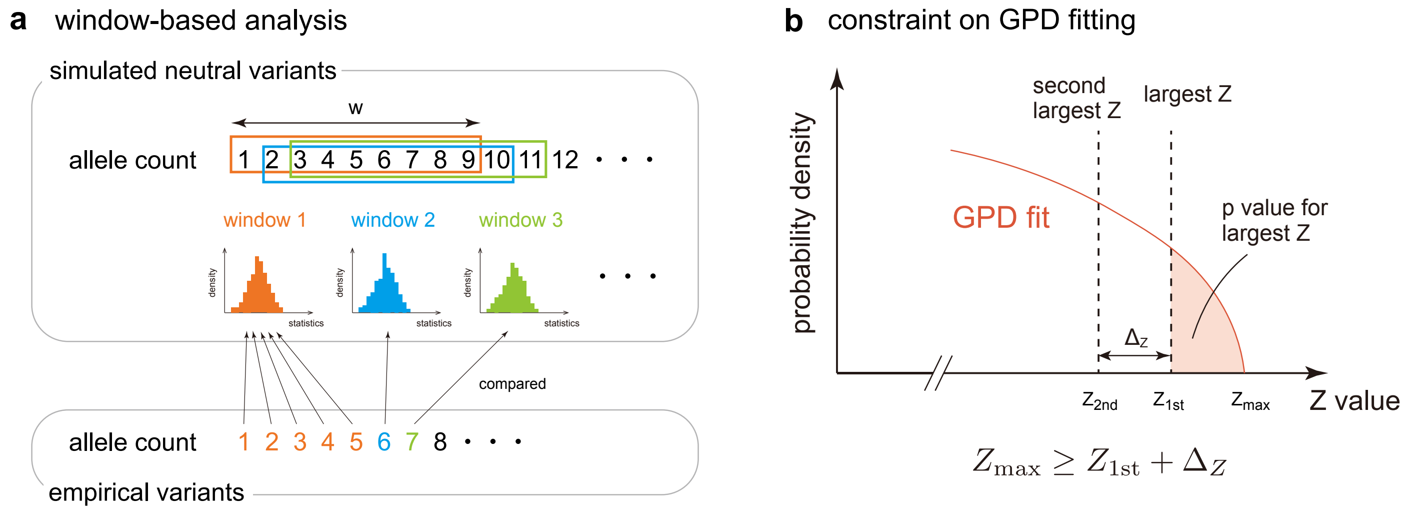


Figure S8: Schematic view of our $p$-value estimation procedure. (a) First, we set a sliding window of minor allele counts with width $w$. Next, we ran simulations to obtain $n_{\text{mut}}$ neutral variants for each minor allele count. In this paper, we used $w = 9$ and $n_{\text{mut}} = 11{,}112$ so that each window spans the frequency range of 0.025 and contains $\sim 10^5$ variants. For each window, the distribution of $Z$-values is smoothed and extended into the extreme tail. Each empirical SNP was matched to the window with the closest mean minor-allele count. (b) In fitting the generalized Pareto distribution, we imposed the constraint that $Z_{1\text{st}} + \Delta_Z$ exist within the support of the fitted distribution, where $\Delta_Z$ is the difference between the largest possible $Z$-value ($Z_{1\text{st}}$) and the second largest ($Z_{2\text{nd}}$). This ensures that the tail probability for $Z_{1\text{st}}$ is integrated over an interval of at least $\Delta_Z$, and prevents unrealistically small $p$-values from being assigned to extreme $Z$-values.

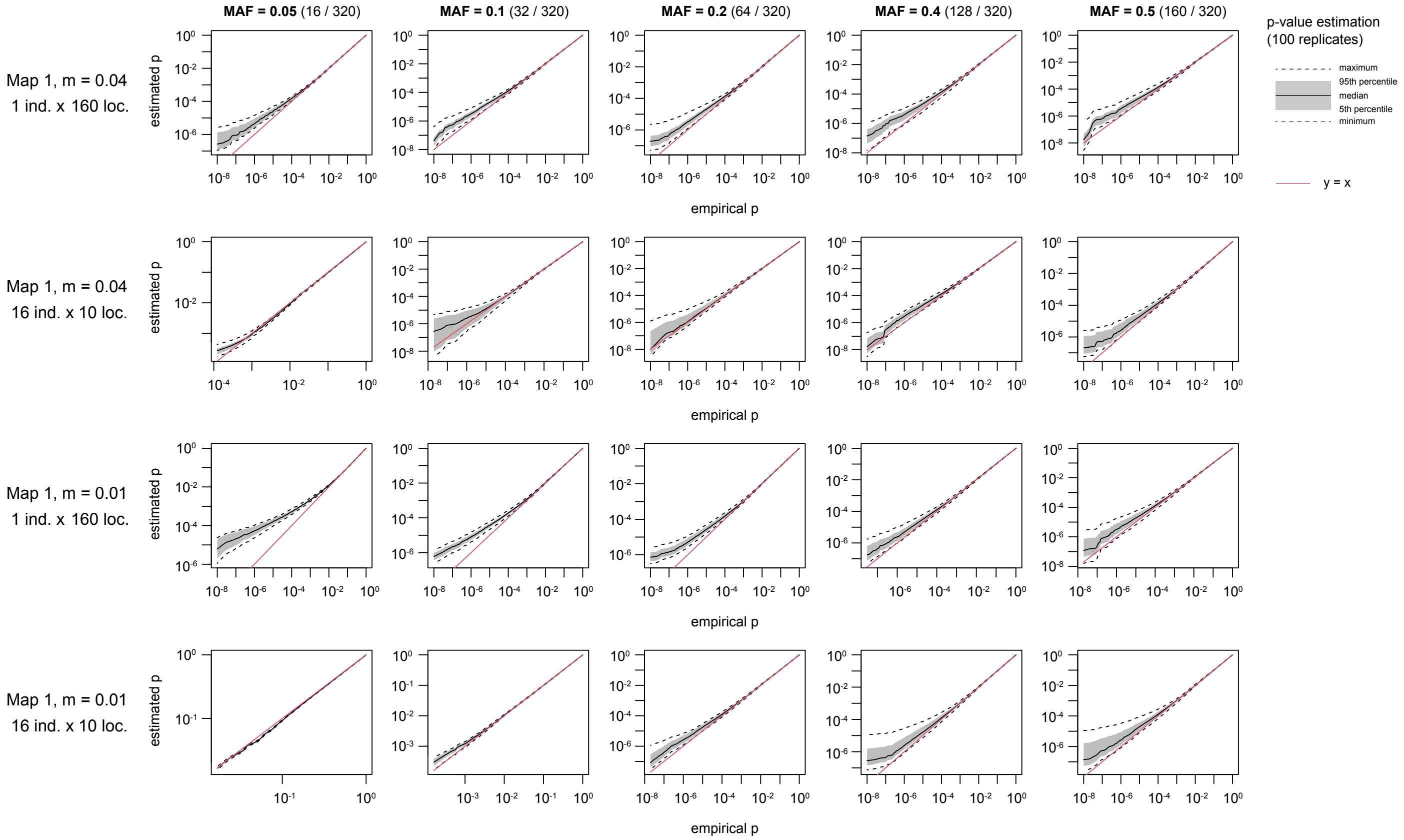


Figure S9: A comparison of the $p$-value estimated using Gaussian kernel smoothing and generalized Pareto fitting from $\sim 10^5$ neutral variants and the empirical $p$-value from $10^8$ variants. The estimation of the $p$-value was conducted 100 times and its percentiles across the runs are reported. Neutral alleles were generated using a migration matrix produced by SimGEA from a 2D-stepping stone linkage-equilibrium dataset with environment map 1.

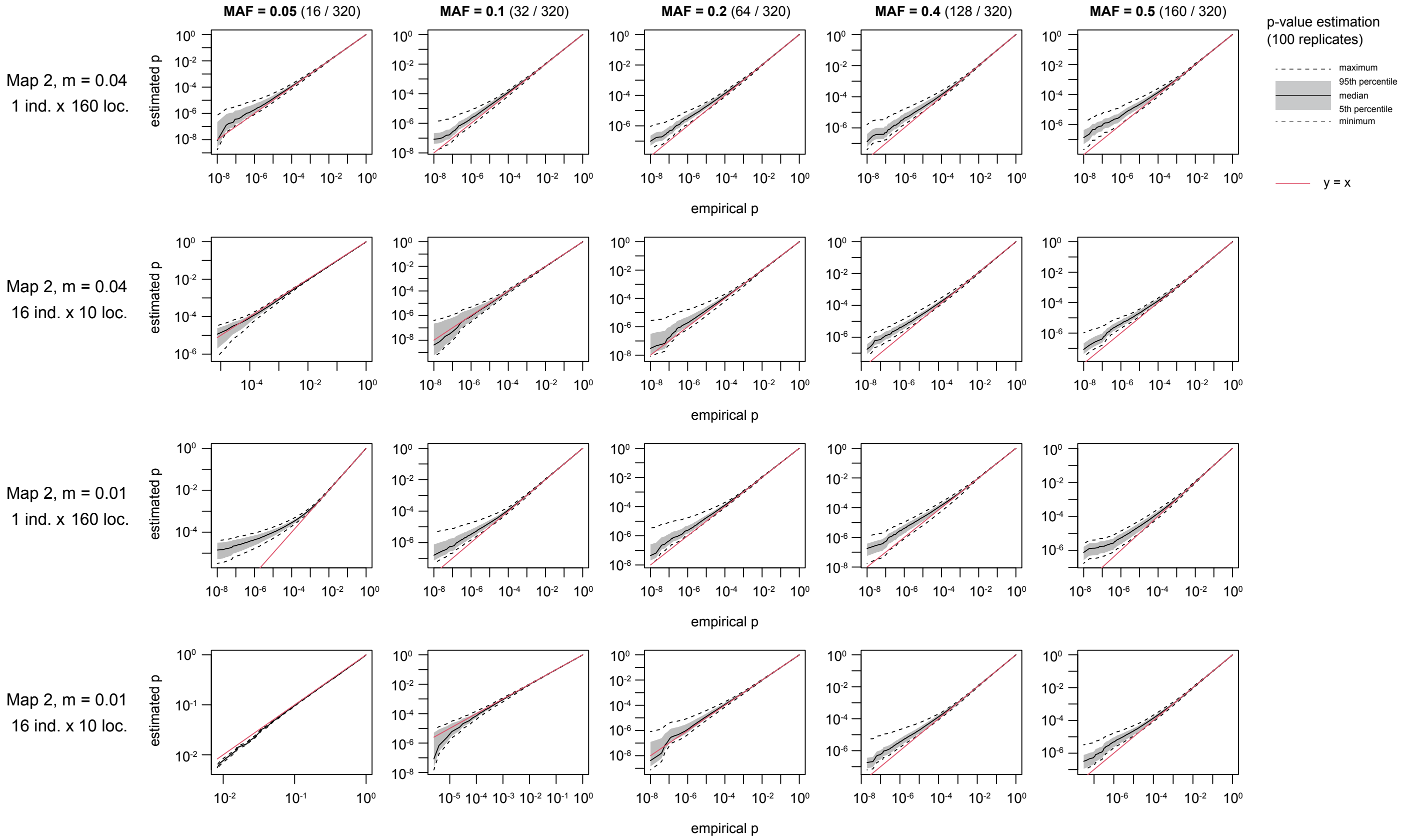


Figure S10: A comparison of the $p$-value estimated using Gaussian kernel smoothing and generalized Pareto fitting from $\sim 10^5$ neutral variants and the empirical $p$-value from $10^8$ variants. The estimation of the $p$-value was conducted 100 times and its percentiles across the runs are reported. Neutral alleles were generated using a migration matrix produced by SimGEA from a 2D-stepping stone linkage-equilibrium dataset with environment map 2.

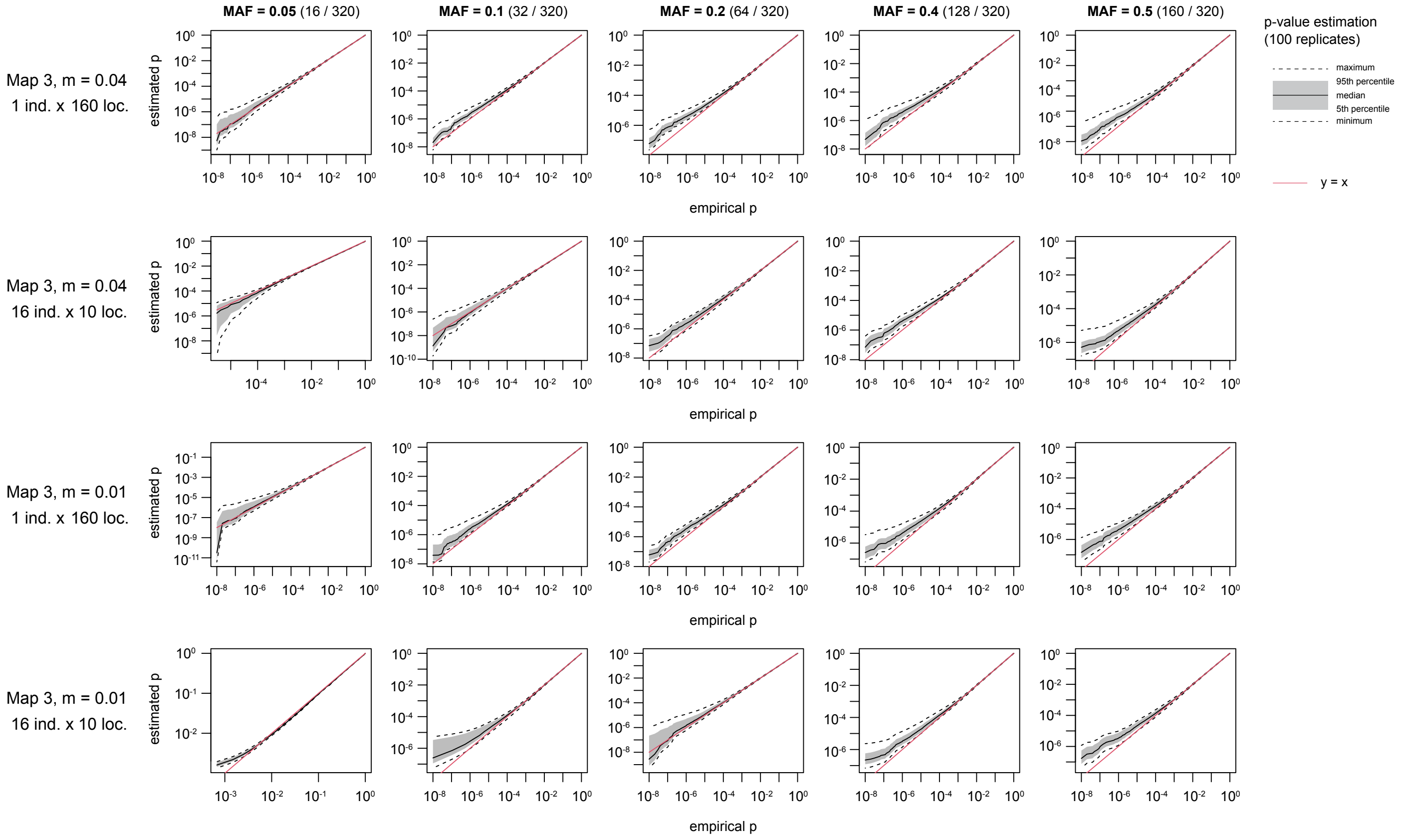


Figure S11: A comparison of the $p$-value estimated using Gaussian kernel smoothing and generalized Pareto fitting from $\sim 10^5$ neutral variants and the empirical $p$-value from $10^8$ variants. The estimation of the $p$-value was conducted 100 times and its percentiles across the runs are reported. Neutral alleles were generated using a migration matrix produced by SimGEA from a 2D-stepping stone linkage-equilibrium dataset with environment map 3.
